# Myeloid deficiency of the intrinsic clock protein BMAL1 accelerates cognitive aging by disrupting microglial synaptic pruning

**DOI:** 10.1101/2022.10.17.512618

**Authors:** Chinyere Agbaegbu Iweka, Erica Seigneur, Amira Latif Hernandez, Sur Herrera Paredes, Mica Cabrera, Eran Blacher, Connie Tsai Pasternak, Frank M. Longo, Luis de Lecea, Katrin I. Andreasson

## Abstract

Aging is associated with loss of circadian immune responses and circadian gene transcription in peripheral macrophages. Microglia, the resident macrophages of the brain, also show diurnal rhythmicity in regulating local immune responses and synaptic remodeling. To investigate the interaction between aging and microglial circadian rhythmicity, we examined mice deficient in the core clock transcription factor, BMAL1. Aging *Cd11b^cre^;Bmal^lox/lox^* mice demonstrated accelerated cognitive decline in association with suppressed hippocampal long-term potentiation and increases in immature dendritic spines. C1q deposition at synapses and synaptic engulfment were significantly decreased in aging *Bmal1*-deficient microglia, suggesting that BMAL1 plays a role in regulating synaptic pruning in aging. In addition to accelerated age-associated hippocampal deficits, *Cd11b^cre^;Bmal^lox/lox^* mice also showed deficits in the sleep-wake cycle with increased wakefulness across light and dark phases. These results highlight an essential role of microglial BMAL1 in maintenance of synapse homeostasis in the aging brain.

**Significance Statement:** This study demonstrates that myeloid deficiency of the circadian clock gene *Bmal1* disrupts microglial synaptic pruning in the hippocampus, accelerates age-associated cognitive decline, and disrupts the sleep-wake cycle.

## Introduction

The circadian clock is a time-keeping system that enables anticipation and adaptation to changes in the environment and maintenance of homeostasis in living organisms (Welz et al., 2019; Patke et al., 2020). Disruption of the circadian clock at the whole-body or organ level is associated with cardiovascular, metabolic, neoplastic and neurological disorders, and is a prominent feature in aging in many organ systems (Yamazaki et al., 2002; Shimizu et al., 2016; Chellappa et al., 2018; Thosar et al., 2018; Wu et al., 2019).

A central regulator of the cellular clock machinery is BMAL1 (brain and muscle Arnt (aryl hydrocarbon receptor nuclear translocator)-like protein) which dimerizes with CLOCK (circadian locomotor output cycles kaput) to regulate circadian gene expression in a cell-specific manner, where approximately 15% of all genes are rhythmically regulated within all cell types (Partch et al., 2014; Trott and Menet, 2018). BMAL1 is a member of the basic-helix–loop–helix (bHLH)-PER–ARNT–SIM (PAS) superfamily of transcription factors and was originally characterized by its high expression in the brain and muscle (Ikeda and Nomura, 1997). Global deletion of *Bmal1* accelerates age-related pathologies such as atherosclerosis, calcification of limb joints, and ocular abnormalities, and reduces lifespan (Kondratov et al., 2006). In peripheral macrophages, *Bmal1* deletion disrupts the diurnal rhythmicity of immune cell trafficking, mitochondrial metabolism and inflammatory responses (Nguyen et al., 2013; Early et al., 2018; Alexander et al., 2020).

Microglia are brain-resident macrophages that maintain tissue homeostasis by clearing cellular debris and misfolded proteins and protecting against pathogens. Microglia also regulate synaptic circuitry by pruning non-functional synapses (Lim et al., 2013; Miyamoto et al., 2016; Wang et al., 2020). Several studies have demonstrated circadian rhythmicity in microglial expression of inflammatory molecules, synaptic pruning and changes in morphology that are altered with age (Hayashi et al., 2013; Fonken et al., 2015; Fonken et al., 2016; Takayama et al., 2016; Nakazato et al., 2017). However, the role of circadian regulation of microglial function in age-associated cognitive decline has not been explored. Here, we investigated the effects of circadian clock disruption in aging microglia by partial ablation of the core clock protein BMAL1 in myeloid-lineage cells. We find that microglia deficient in BMAL1 show disrupted synaptic pruning and a persistence of immature dendritic spines, a phenotype associated with accelerated age-related hippocampal memory decline and disrupted sleep-wake cycle.

## Results

### Microglial BMAL1 deficiency disrupts hippocampal-dependent memory in aged mice

The circadian clock declines with aging and is associated with deficits in learning and memory (Fonseca Costa and Ripperger, 2015; Chellappa et al., 2018; Adler et al., 2019). We have previously shown that macrophage circadian gene expression and immune responses decline in aging mice (Blacher et al., 2021). Aging microglia also exhibit a diminished capacity for normal function that is associated with impaired cognition (Harry, 2013; Kaneshwaran et al., 2019; Duggan and Parikh, 2021). To investigate the interaction between microglial circadian function and aging, we generated myeloid-specific *Bmal1* conditional knock out mice (*CD11b^cre^;Bmal1^lox/lox^*; cKO) and confirmed reduced BMAL1 protein expression in peritoneal macrophages (**Fig. S1 A**; *p=0.0178*). We then subjected young and aged WT and cKO mice to a series of cognitive tasks. The novel object recognition task, a non-spatial recall test that measures discrimination between a familiar and novel object, revealed impaired memory in aged *Bmal1* cKO compared to littermate WT mice (**Fig. 1 A**). Young *Bmal1* cKO mice performed similarly to WT in their examination of the novel object (**Fig. 1 A**). Spatial learning and memory, which is encoded primarily by the hippocampus, was tested using the Barnes maze task. Primary escape latency over the 5-day training period decreased by a larger degree for young WT compared to aged WT and *Bmal1* cKO mice (**Fig. 1 B**). Aged *Bmal1* cKO showed a significant increase in escape latency compared to WT mice (**Fig. 1 C**; *p=0.0118*), indicating a deficit in spatial memory. Young *Bmal1* cKO performed better on the Barnes maze task compared to WT mice (**Fig. 1 C**; *p=0.0319*) consistent with a recent study showing that microglial loss of *Bmal1* improved cognition in young mice (Wang et al., 2021). In the open field test, which assesses motor function, exploratory activity, and anxiety, there were no differences in motor function but there was increased anxiety in aged *Bmal1* cKO compared to WT mice (**Fig. 1 D-E**; *p=0.0301*). There were no differences in motor activity, exploration or anxiety between young *Bmal1* cKO and WT mice (**Fig. 1 D-E**). These results indicate that microglial BMAL1 deficiency accelerates age-associated deficits in spatial and non-spatial learning and memory.

**Figure 1.**
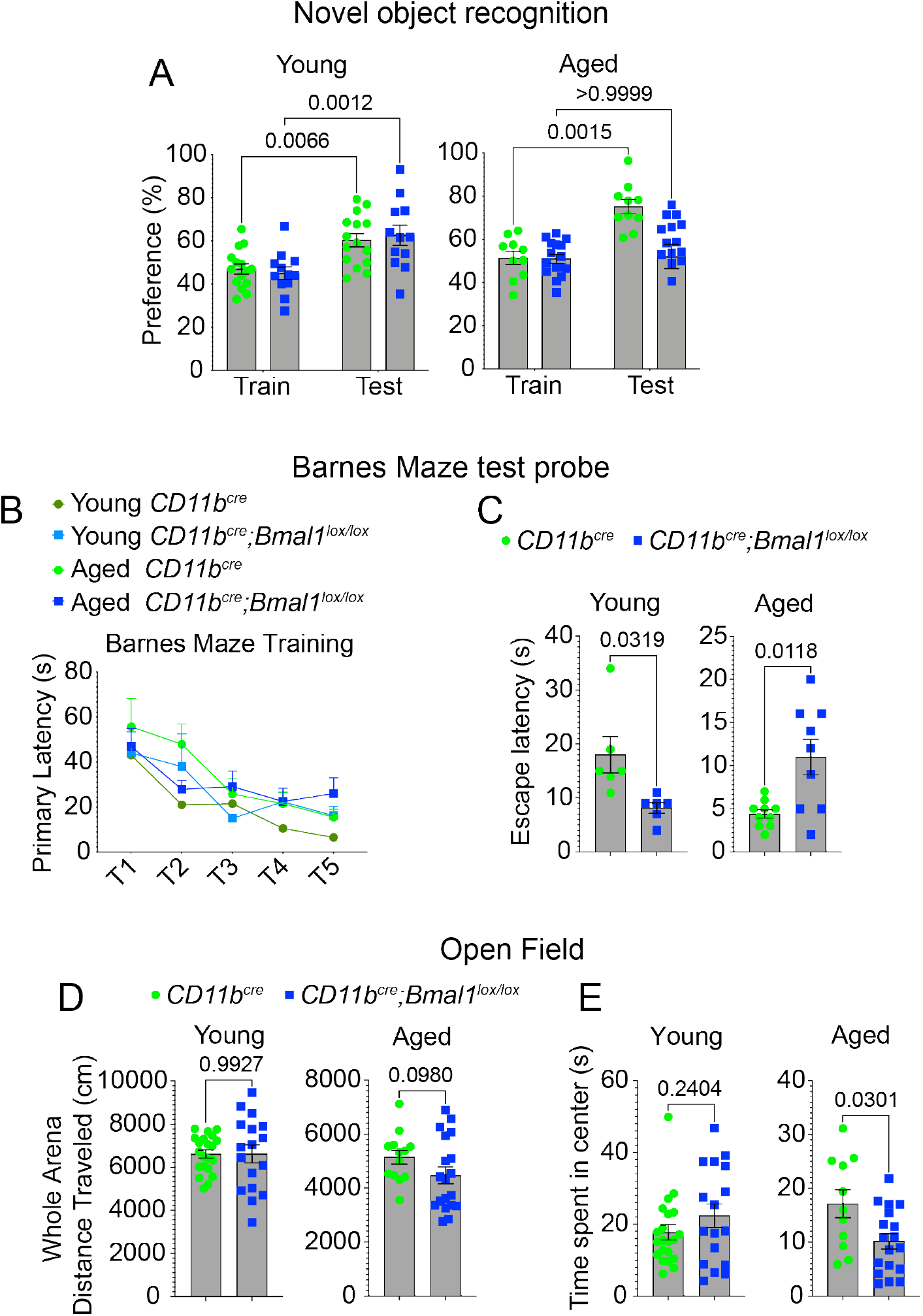
Microglia BMAL1 deficiency disrupts hippocampal-dependent behavior and increases anxiety in aged, but not young mice. **(A)** Preference in the novel object recognition task in young (3-6 months, *CD11b^cre^*, n = 15; *CD11b^cre^;Bmal1*^*lox*/lox^, n = 12) and aged (18-20 months, *CD11b^cre^*, n = 10; *CD11b^cre^;Bmal1^lox/lox^*, n = 16) mice. **(B)** Primary escape latency during the 5-day trial period for young and aged mice. **(C)** Escape latency in the testing phase of the Barnes maze task in young (3-6 months, *CD11b^cre^*, n = 6; *CD11b^cre^;Bmal1^lox/lox^*, n = 6) and aged (18-20 months, *CD11b^cre^*, n = 9; *CD11b^cre^;Bmal1^lox/lox^*, n = 10) mice. **(D)** Distance travelled in whole arena in the open field task in young (3-6 months, *CD11b^cre^*, n = 21; *CD11b^cre^;Bmal1^lox/lox^*, n = 17) and aged (18-20 months, *CD11b^cre^*, n = 12; *CD11b^cre^;Bmal1^lox/lox^*, n = 19) mice. **(E)** Time spent in the center area in the open field task in young (3-6 months, *CD11b^cre^*, n = 21; *CD11b^cre^;Bmal1^lox/lox^*, n = 17) and aged (18-20 months, *CD11b^cre^*, n = 12; *CD11b^cre^;Bmal1^lox/lox^*, n = 19) mice. Data are represented as the mean ± SEM. *P*-values were calculated using paired *t*-test, or two-tailed Student’s *t*-test.

### Microglial BMAL1 deficiency suppresses long-term potentiation in the CA1 hippocampal region of aged mice

Next, we asked if neural correlates of learning and memory in aged mice were altered in *Bmal1* cKO mice. First, we evaluated short-term synaptic plasticity by measuring paired-pulse ratio (PPR) in the CA1 region of the hippocampus. PPR measures the probability of activity-dependent pre-synaptic vesicular release following an action potential (Katz and Miledi, 1968). The ratio of the amplitude of the second response to that of the first to stimulation is inversely related to the release probability. PPR was significantly reduced at 10 ms (**Fig. 2 A**; *p=0.0339*) and increased at 50 ms (*p=0.0049*), 100 ms (*p=0.0595*), 200 ms (*p=0.0467*), and 500 ms (*p=0.0404*) in aged *Bmal1* cKO compared to WT mice (**Fig. 2 A**), consistent with perturbation of pre-synaptic short-term plasticity. The input/output (I/O) curve reflecting the efficacy of pre- and post-synaptic neurotransmitter release was significantly increased at higher stimulation strengths (50 to 65 μA) in aged *Bmal1* cKO mice, consistent with disrupted basal synaptic transmission (**Fig. 2 B**; *p=0.0231*).

**Figure 2.**
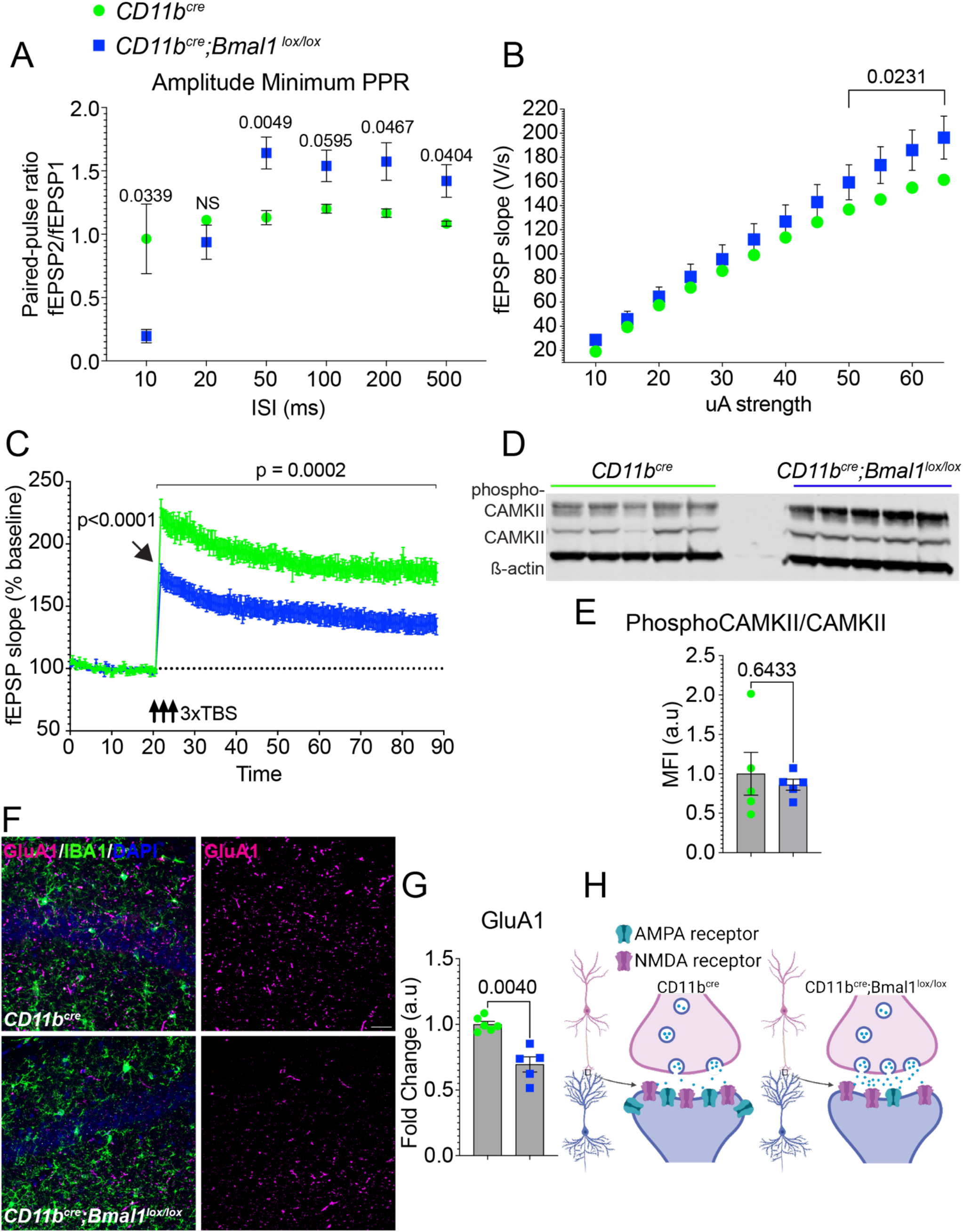
Deficits in long-term potentiation (LTP) in the CA1 region of the hippocampus in aged *Bmal1* cKO mice. **(A)** Paired-pulse ratio was recorded from CA1 pyramidal neurons with inter-stimulus intervals: 10, 20, 50, 100, 200, or 500 ms from aged *CD11b^cre^* (18-20 months; n = 8) and *CD11b^cre^;Bmal1^lox/lox^* (n = 10) mice. **(B)** Input/output (I/O) curves as a measure of basal synaptic transmission in the CA1 region of the hippocampus (10 slices, 5 mice per aged *CD11b^cre^* and *CD11b^cre^;Bmal1^lox/lox^* mice). **(C)** Long-term potentiation (LTP) in the CA1 hippocampal region over a 90-min recording interval (n = 10 slices, 5 mice per aged *CD11b^cre^* and *CD11b^cre^;Bmal1^lox/lox^* mice). Arrow shows that induction of LTP is significantly reduced in *CD11b^cre^* (228.669 ± 7.285) and *CD11b^cre^;Bmal1^lox/lox^* (176.359 ± 7.593; *p<0.0001*). **(D)** Representative immunoblot of phospho-CAMKII and CAMKII in aged *CD11b^cre^* and *CD11b^cre^;Bmal1^lox/lox^* mice (18-20 months; n = 5/group). **(E)** Quantification of phospho-CAMKII and CAMKII immunoblot in **(D)** normalized to ß-actin. **(F)** Representative confocal images of GluA1 (magenta) expression in the hippocampal CA1 region of aged *CD11b^cre^* (top, n = 6) and *CD11b^cre^;Bmal1^lox/lox^* (bottom, n = 5) mice. Scale bar, 50 μm. **(G)** Quantification of mean fluorescence intensity of GluA1 in the hippocampal CA1 region of aged *CD11b^cre^* (n = 6) and *CD11b^cre^;Bmal1^lox/lox^* (n = 5) mice. **(H)** Diagram highlighting differences in synaptic release and numbers of receptors between genotypes in aged CA1 hippocampus. Data are represented as the mean ± SEM. *P*-values were calculated using Two-way RM ANOVA or Two-way ANOVA: effects of time and genotype and Sidak’s multiple comparisons test with Geisser-Greenhouse correction. Theta burst stimulation (TBS).

We next examined long-lasting, activity-dependent changes in synaptic efficacy from the CA3 to CA1 Schaffer collateral pathway in the hippocampus to assess post-synaptic plasticity. We found significant impairment in long-term potentiation (LTP) in hippocampal slices from aged *Bmal1* cKO compared to WT mice across 70 minutes of recording (Fig. 2 C; main effect of genotype for 70 minutes post-induction: F_1,18_= 20.77, *p=0.0002*). This impairment was apparent immediately after LTP induction (**Fig. 2 C**; *p<0.0001*). Thereafter, LTP in aged *Bmal1* cKO mice slowly decayed to ~35% potentiation, whereas WT mice stabilized LTP at ~80% with respect to baseline.

Calcium/calmodulin-dependent protein kinase II (CaMKII) is critical for the induction of LTP, and inhibition of CAMKII prevents LTP (Leonard et al., 1999; Barria and Malinow, 2005; Lee et al., 2009; Lisman et al., 2012). To investigate the candidate mechanisms underlying the reduced LTP in aged *Bmal1* cKO mice, we tested whether microglial *Bmal1* deficiency might alter hippocampal levels of CAMKII or phosphorylated CAMKII. Quantitative immunoblotting showed increased total CAMKII expression and CAMKII phosphorylation, however, there was no difference in the ratio of phosphorylated CAMKII/total CAMKII between aged *Bmal1* cKO and WT mice (**Fig. 2 D-E**; *p=0.6433*). The AMPA receptor subunit, GluA1, mediates LTP maintenance by enhancing AMPA receptor conductance and anchoring at the synapse (Jiang et al., 2021). To reconcile the increase in I/O ratio with decreased LTP induction and amplitude, we assessed levels of GluA1 expression. Aged *Bmal1* cKO mice exhibited decreased GluA1 signal in the CA1 hippocampal region compared to WT mice (**Fig. 2 F-H**; *p=0.0040*). These data suggest that microglial BMAL1 deficiency alters hippocampal plasticity in aged mice by a mechanism involving reduced insertion of AMPA receptors in dendritic spines in aged mice.

### BMAL1-deficient microglia increase synaptic density in the CA1 hippocampal region of aged mice

Microglia actively maintain neural circuitry and plasticity through selective pruning of synapses (Schafer and Stevens, 2010; Paolicelli et al., 2011; Schafer et al., 2012). Given the cognitive deficits and altered synaptic plasticity observed in aged *Bmal1* cKO mice, we measured expression levels of presynaptic and postsynaptic proteins, SNAP25 and PSD95, respectively, in the CA1 hippocampal region. Aged *Bmal1* cKO mice showed higher immunoreactivity for both PSD95 (*p=0.0487*) and SNAP25 (*p=0.0439*) compared to aged WT mice (**Fig. 3 A-B**). No significant differences were observed between young *Bmal1* cKO and WT mice (**Fig. S1 B-C**). These results were confirmed using quantitative immunoblotting for PSD95 (*p=0.0010*) and SNAP25 (*p=0.0441*) (**Fig. 3 C-D**). Assessment of synaptic morphology and density by Golgi staining confirmed increased dendritic spine density in aged *Bmal1* cKO compared to WT mice (**Fig. 3 E-F**; *p=0.0245*). Aged *Bmal1* cKO mice had a higher density of filopodia-like (*p=0.00497*), stubby (*p=0.0185*) and mushroom spines (*p=0.01498*) compared to WT mice (**Fig. 3 G**), indicating a persistence of immature synapses in aged *Bmal1* cKO mice. These data suggest a significant impairment in pruning of synapses resulting from microglial BMAL1 deficiency in aged mice.

**Figure 3.**
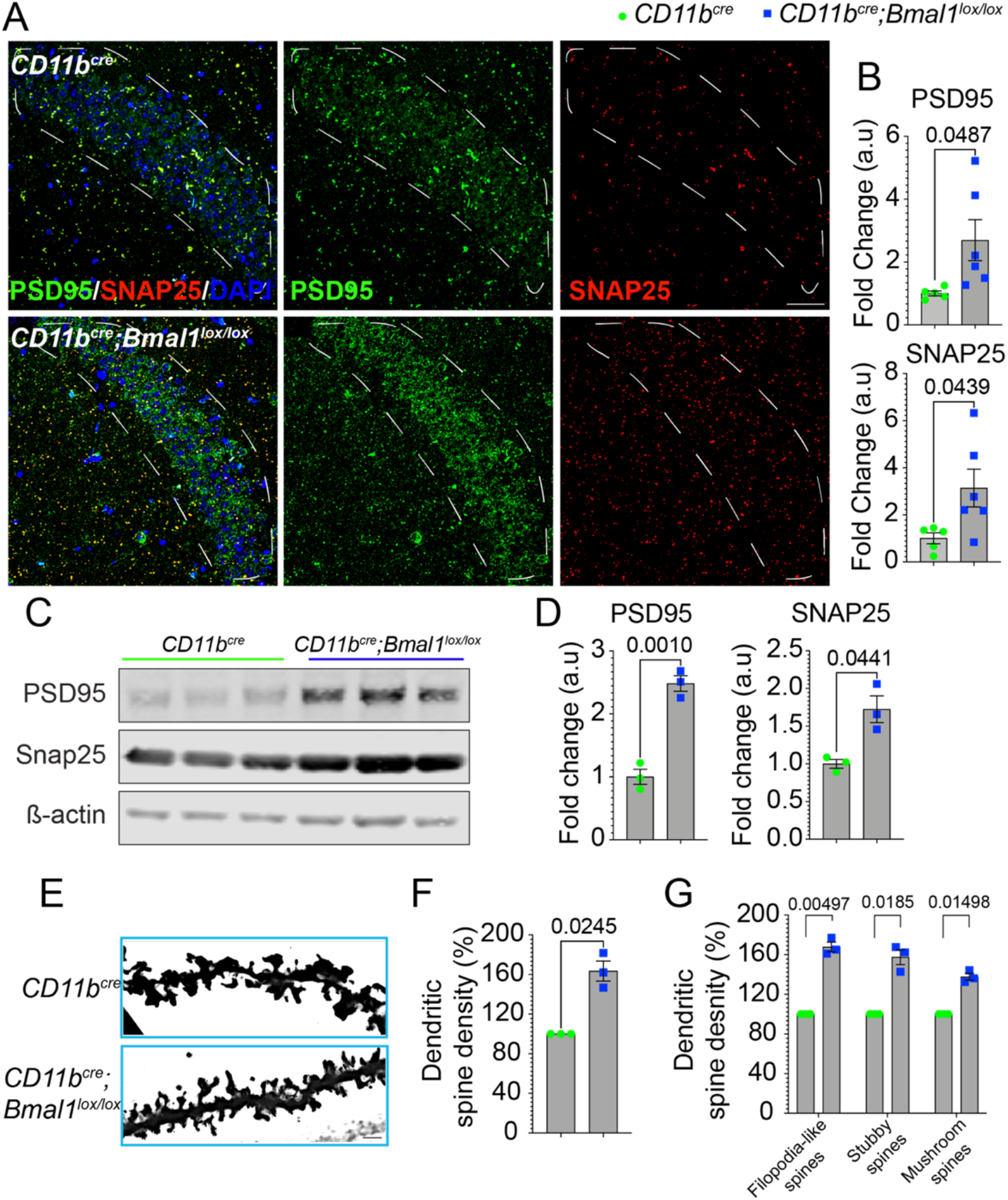
Increased numbers of immature spines in aged *Bmal* cKO CA1 hippocampus. **(A)** Representative images of PSD95 (green) and SNAP25 (red) expression in aged *CD11b^cre^* and *CD11b^cre^;Bmal1^lox/lox^* mice (18-20 months). Scale bar, 50 μm. White dotted lines in **(A)** indicate region of interest quantified in **(B)**. **(B)** Mean fluorescence intensity of PSD95 and SNAP25 in aged *CD11b^cre^* (n = 5) and *CD11b^cre^;Bmal1^lox/lox^* mice (n = 6). **(C)** Representative immunoblot of PSD95 and SNAP25 in aged *CD11b^cre^* and *CD11b^cre^;Bmal1^lox/lox^* mice (18-20 months; n = 3/group). **(D)** Quantification of PSD95 and SNAP25 normalized to ß-actin. **(E)** Representative images of Golgi stain of apical dendritic spines in the CA1 region of the hippocampus aged *CD11b^cre^* and *CD11b^cre^;Bmal1^lox/lox^* mice (18-20 months; n = 3/group). Scale bar, 5 μm. **(F)** Dendritic spine density (10-12 neurons/CA1 region) in aged *CD11b^cre^* and *CD11b^cre^;Bmal1^lox/lox^* (n = 3/group) mice. **(G)** Dendritic spine density of filopodia-like, stubby and mushroom spines in aged *CD11b^cre^* and *CD11b^cre^;Bmal1^lox/lox^* (n = 3/group) mice. Data are represented as the mean ± SEM. *P*-values were calculated using two-tailed Student’s *t*-test.

### Microglial BMAL1 deficiency decreases C1q expression and synaptic pruning in aged CA1 hippocampus

To investigate mechanisms that may underlie defective synaptic pruning in aged *Bmal1* cKO mice, we measured expression of C1q, an opsonin that is deposited on synaptic terminals and promotes synapse removal through interaction with the microglial C3 receptor, CR3 (Stevens et al., 2007; Schafer et al., 2012; Hong et al., 2016). C1q is primarily expressed by microglia and increases with age, although neuronal expression has also been reported (Stevens et al., 2007; Stephan et al., 2013; Fonseca et al., 2017). In the CA1 hippocampal region of aged *Bmal1* cKO mice, C1q colocalized with PSD95 but its abundance was significantly reduced (**Fig. 4 A-B**; *p=0.0010*) in contrast to PSD95 levels (**Fig. 4 A-B**; *p=0.0012*) which were significantly elevated (**Fig. 3 A-B**). CD68, a microglial lysosomal protein, was observed in close proximity to PSD95/C1q and was also decreased in aged *Bmal1* cKO (**Fig. 4 A-B**; *p=0.0427*), suggesting reduced microglial production of both C1q and CD68. There were no differences in C1q, PSD95 or CD68 levels between young *Bmal1* cKO and WT mice (**Fig. S2 A-B**).

**Figure 4.**
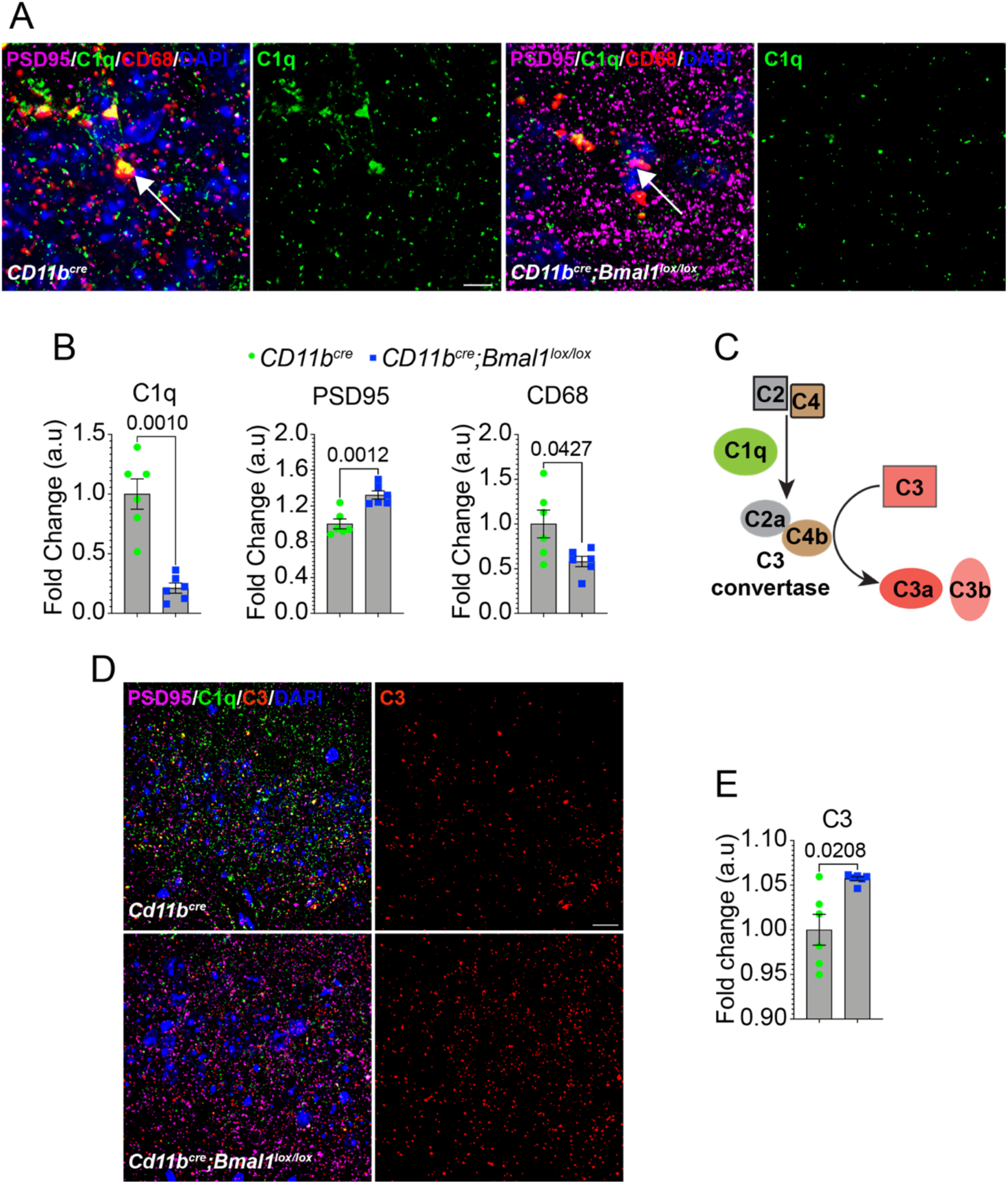
Microglial BMAL1 deficiency decreases C1q and increases C3 in aged CA1 hippocampus. **(A)** Representative images of C1q (green), PSD95 (magenta) and CD68 (red) expression in the hippocampal CA1 region of aged *CD11b^cre^* and *CD11b^cre^;Bmal1^lox/lox^* mice (18-20 months). Scale bar, 10 μm. White arrow shows colocalization of C1q, PSD95 and CD68 present in aged *CD11b^cre^* that is absent in *CD11b^cre^;Bmal1^lox/lox^* mice. **(B)** Mean fluorescence intensity (MFI) of C1q, PSD95 and CD68 (n = 6/group). **(C)** Schematic illustrating C1q activation of C3. **(D)** Representative confocal images of C1q (green), PSD95 (magenta) and C3 (red) expression in the CA1 hippocampal area in aged *CD11b^cre^* and *CD11b^cre^;Bmal1^lox/lox^* mice. Scale bar, 20 μm. **(E)** MFI of C3 in aged *CD11b^cre^* and *CD11b^cre^;Bmal1^lox/lox^* mice (n = 6/group). Data are represented as the mean ± SEM. *P*-values were calculated using two-tailed Student’s *t*-test.

C1q promotes the formation of C3 convertase through activation of complement proteins, C4 and C2. C3 convertase cleaves C3 to produce C3a, a pro-inflammatory mediator, and C3b, an opsonin that initiates phagocytosis via the CR3 receptor on microglia (Presumey et al., 2017). We observed a reciprocal increase in C3 immunoreactivity in the CA1 hippocampal region of aged *Bmal1* cKO mice compared to WT (**Fig. 4 C-E**; *p=0.0208*), suggesting that C3 may accumulate in response to reduced levels of C3 convertase.

Next, we assessed microglial engulfment of synapses by measuring colocalization of PSD95 and CD68 within IBA1+ microglia. Superresolution microscopy was used to quantify PSD95 within CD68+ structures in IBA+ microglia in order to identify engulfed synapses. Aged *Bmal1* cKO microglia demonstrated a significant decrease in internalized PSD95 compared to WT (**Fig. 5 A-C**; *p=0.0469*; **Video 1**), consistent with decreased synapse engulfment. Taken together, these data indicate that microglial loss of BMAL1 decreases C1q deposition on synapses and synaptic engulfment.

**Figure 5.**
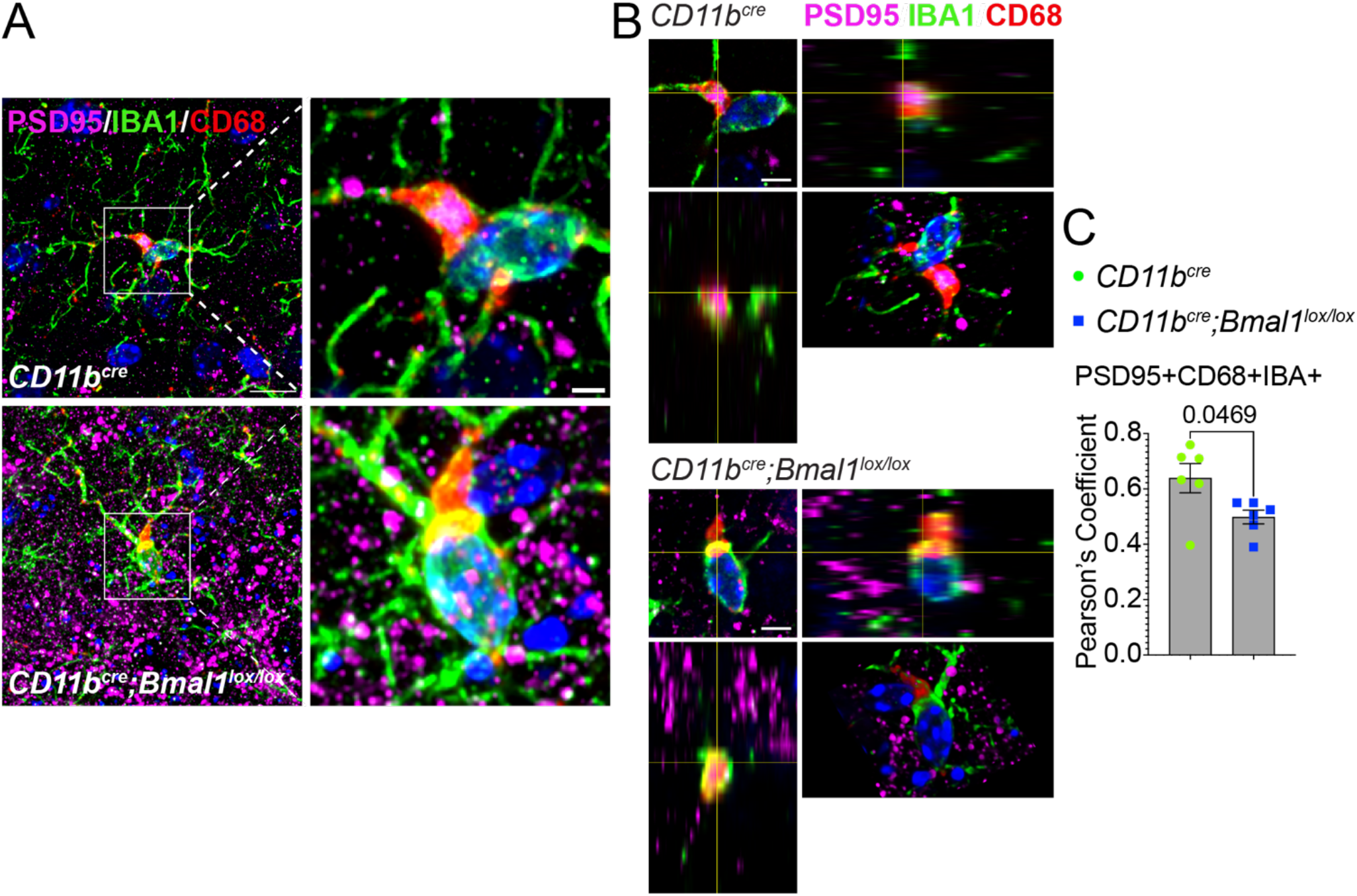
Microglial BMAL1 deficiency decreases synaptic engulfment in aged mice. **(A)** Representative images of IBA1+/CD68+ cell engulfment of PSD95+ particles in microglia in aged *CD11b^cre^ and CD11b^cre^;Bmal1^lox/lox^* mice. Scale bar, 2 μm. **(B)** Orthogonal views and rotated 3D projection of PSD95+ particles in IBA+/CD68+ microglia from aged *CD11b^cre^ and CD11b^cre^;Bmal1^lox/lox^* mice. **(C)** Pearson’s coefficient of colocalization of PSD95 and CD68 in IBA1+ cells (n = 6/group) mice. Data are represented as the mean ± SEM. *P*-values were calculated using two-tailed Student’s *t*-test.

### Microglia deficient in BMAL1 exhibit microgliosis and reduced expression of lysosomal proteins in the CA1 hippocampal region

To further characterize *Bmal1*-deficient microglia, we measured IBA1 and CD68 expression in the CA1 hippocampal region. A significant increase was observed in IBA1 immunoreactivity (*p=0.0028*) and number of microglia (*p=0.0126*) in aged *Bmal1* cKO compared to WT mice (**Fig. 6 A-C**). Quantitative immunoblot analysis confirmed the increase in IBA1 expression in aged *Bmal1* cKO mice (**Fig. 6 D-E**; *p=0.0067*). Consistent with our previous observation shown in **Fig. 4 B**, CD68 was significantly decreased in aged *Bmal1* cKO mice (**Fig. 6 A-C**; *p=0.0494*). The proportion of IBA1+ cells immunoreactive for CD68 was also significantly reduced, indicating an overall decrease in CD68 expression (**Fig. 6 A-C**; *p=0.0197*). Morphological analysis of microglial complexity revealed decreases in branch length (*p=0.0489*), branch number (*p=0.0385*) and number of junctions (*p=0.0398*) in aged *Bmal1* cKO compared to WT mice (**Fig. 6 F-G**). No differences between young WT and *Bmal1* cKO mice in either IBA1 or CD68 expression, proportion of IBA1+ cells immunoreactive for CD68, number of microglia, or morphological parameters were observed (**Fig. S3 A-E**).

**Figure 6.**
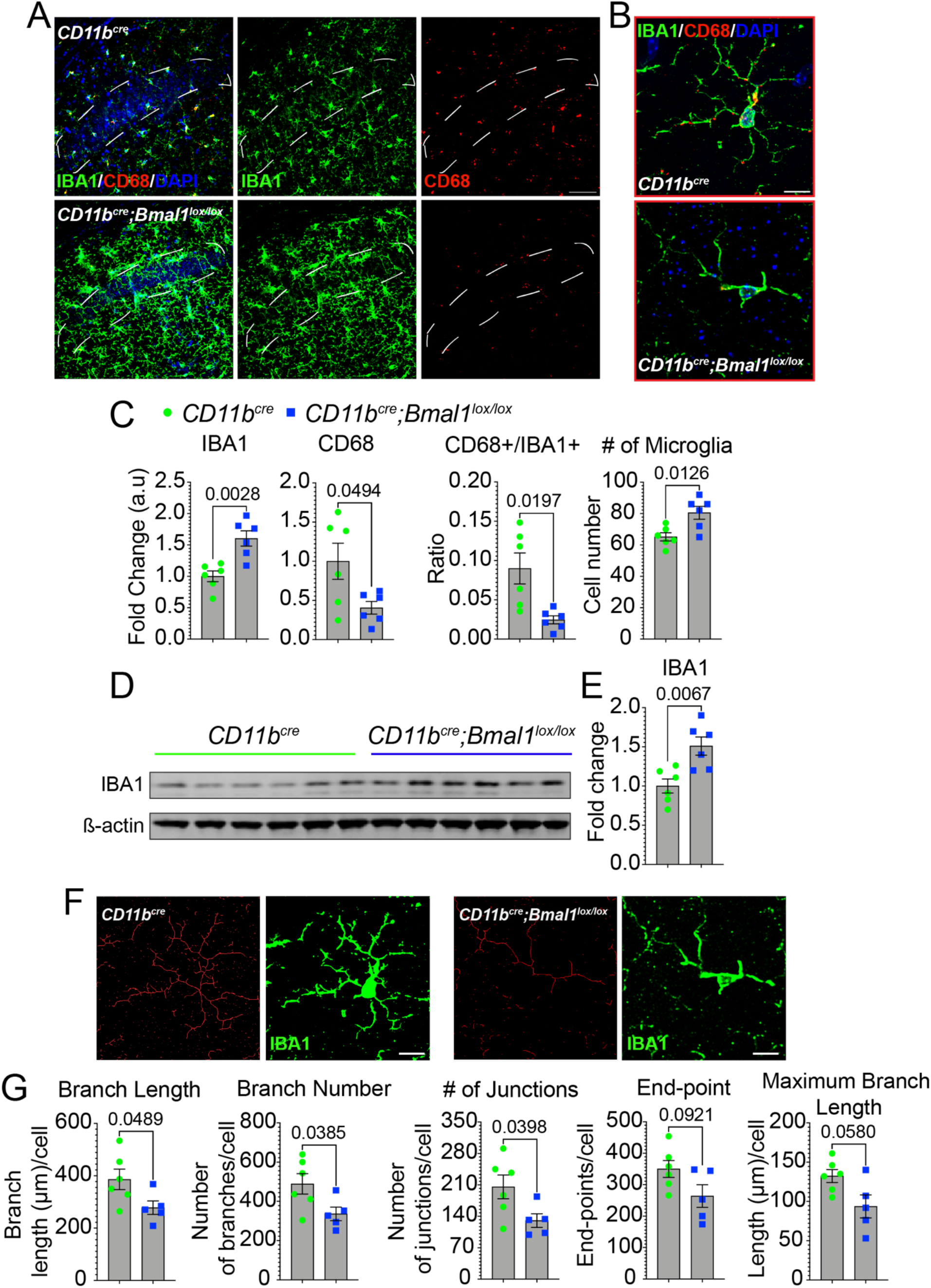
Myeloid *Bmal1* deletion increases microglial activation but decreases lysosomal function in aged mice. **(A)** Representative confocal images of IBA1 (green) and CD68 (red) expression in the hippocampal CA1 region of aged *CD11b^cre^* and *CD11b^cre^;Bmal1^lox/lox^* mice (18-20 months, n = 6/group). Scale bar, 50 μm. White dotted lines in (**A**) indicate region of interest quantified in **(C)**. **(B)** Higher magnification of microglia in CA1 hippocampal region from aged *CD11b^cre^* and *CD11b^cre^;Bmal1^lox/lox^* mice.Scale bar, 5 μm. **(C)** Mean fluorescence intensity of IBA1, CD68, proportion of IBA1-positive (+) cells that are CD68+, and the number of DAPI+IBA1+ microglia (n = 6/group). **(D)** Representative immunoblot of IBA1 in aged *CD11b^cre^* and *CD11b^cre^;Bmal1^lox/lox^* mice (18-20 months; n = 6/group). **(E)** Quantification of IBA1 immunoblot normalized to ß-actin. **(F)** Representative images of skeletonized microglia overlaid on original image from aged *CD11b^cre^* and *CD11b^cre^;Bmal1^lox/lox^* mice. **(G)** Quantification of microglial complexity.Scale bar, 5 μm. Every measurement that contained ≤ 2 endpoints with a maximum branch length of less than the cutoff value of 0.5 μm was removed from the analysis. Data are represented as the mean ± SEM. *P*-values were calculated using two-tailed Student’s *t*-test.

CD68 is a member of the lysosomal/endosomal-associated transmembrane glycoprotein (LAMP) family whose expression is upregulated with inflammatory stimuli such as bacterial lipopolysaccharide (LPS). Microglial CD68 expression and lysosomal dysfunction increases with aging and is associated with neurodegeneration (Wong et al., 2005; Chistiakov et al., 2017; Peng et al., 2019; Root et al., 2021). We investigated additional lysosomal proteins including LAMP1, whose expression was similarly reduced in the CA1 hippocampal region of aged *Bmal1* cKO mice (**Fig. S4 A-B**; *p=0.0335*), as was p62 (**Fig. S4 C-D**; *p=0.0483*) which is required for the formation of the omegasome, a precursor structure of the autophagolysosome (Cha-Molstad et al., 2017; Hsieh and Yang, 2019). These data suggest that although microglia in aged *Bmal1* cKO mice are proliferating and are morphologically activated, their reduced levels of CD68, LAMP1 and p62 suggest a deficit in lysosomal function.

### Deficiency of BMAL1 in microglia alters the omegasome and viral response genes

To identify biological pathways affected by microglial loss of BMAL1, we performed RNA sequencing on CD11b+ microglia isolated from young and aged, WT and *Bmal1* cKO mice. Principal component analysis (PCA) showed a large difference in transcriptional profiles driven by age (PC1) with variability among the aged mice (PC2) (**Fig. 7 A**). No distinction between genotypes in either aged group was observed at the full transcriptome level. At the gene level, differential expression analysis (DEA) revealed 150 differentially expressed genes (DEGs) between *Bmal1* cKO and WT aged mice. 4/19 (21%) down-regulated and 1/144 (<1%) upregulated genes overlapped between young and aged *Bmal1* cKO mice (**Fig. 7 B**). Of those, 13/150 (9%) were down-regulated and 137/150 (91%) were upregulated (**Fig. 7 C**). 18 DEGs were identified between young *Bmal1* cKO and WT mice, with 10/18 (56%) up-regulated and 8/18 (44%) down-regulated genes in *Bmal1* cKO (**Fig. S5**). Gene ontology (GO) analysis showed significant enrichment only in downregulated genes for biological functions that include innate immunity, autophagy and viral response (**Table 1**). Interestingly, GO enrichment for cellular component was significant for the omegasome pathway, in line with our observations of decreased p62 protein levels in aged *Bmal1* cKO microglia.

**Figure 7.**
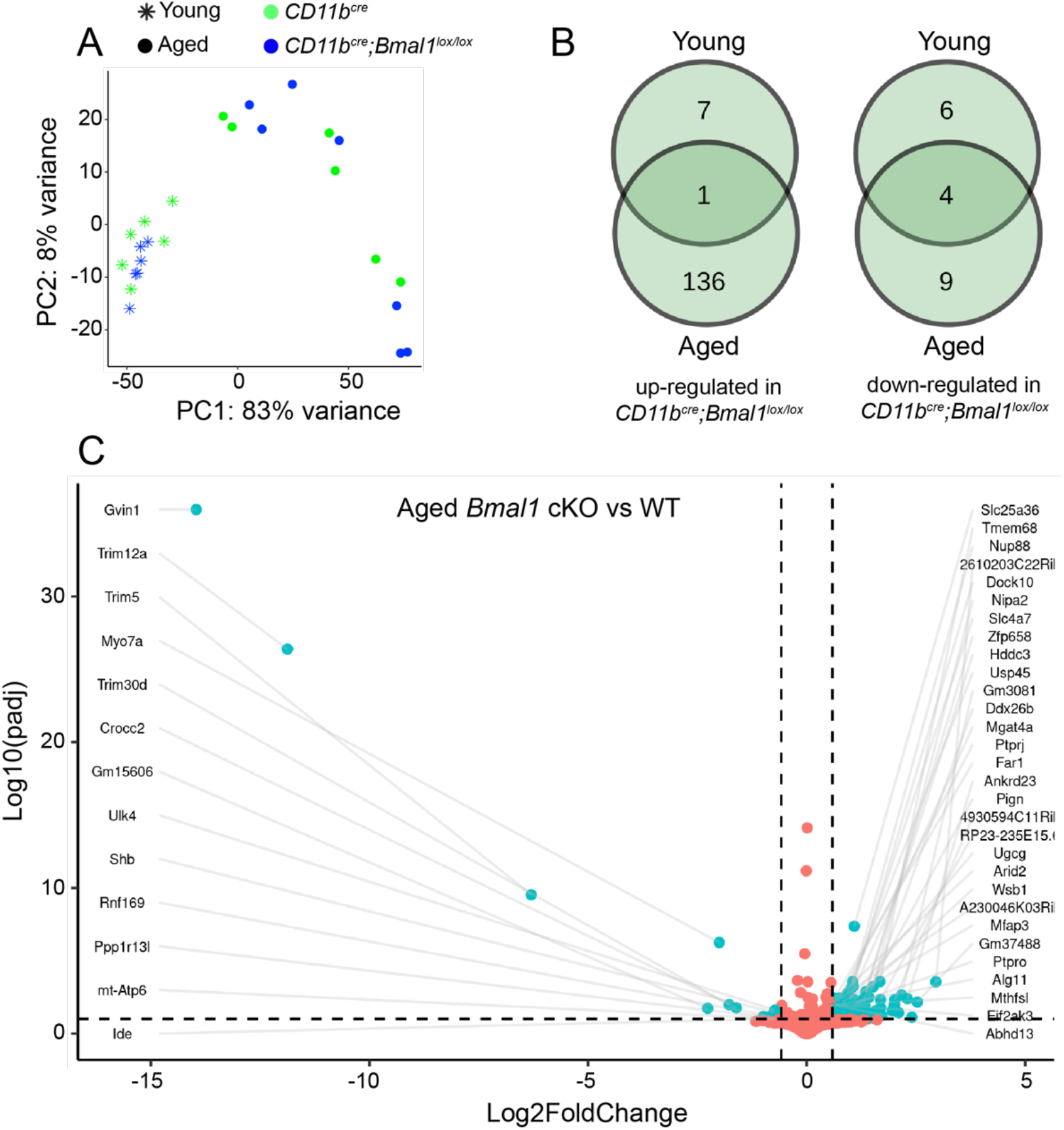
Microglial gene expression changes in young and aged WT and *Bmal1* cKO mice. **(A)** Principal component analysis (PCA) shows distinct clustering of young *CD11b^cre^* and *CD11b^cre^;Bmal1^lox/lox^* (n = 6/group) mice versus aged *CD11b^cre^* (n = 6) and *CD11b^cre^;Bmal1^lox/lox^* (n = 7) mice. **(B)** Venn diagram of number of differentially expressed genes in young and aged *CD11b^cre^;Bmal1^lox/lox^* mice. **(C)** Volcano plot of −log10 p-adjusted value versus −log2 fold change of normalized counts between aged *CD11b^cre^* and *CD11b^cre^;Bmal1^lox/lox^* mice. Each dot represents a single transcript. Cyan dots denote significantly differentially expressed genes. Orange dots denote genes that are unchanged between the genotypes (adjusted p-value <0.05).

### Microglial deficiency of BMAL1 disrupts the sleep-wake cycle

Sleep is a major regulator of microglial function, and neuron-microglia interactions change during wake, NREM and REM states, reflecting changes in synapse remodeling (Deurveilher et al., 2021). Indeed, the sleep-wake cycle drives changes in synaptic ultrastructure and the daily dynamics of synaptic protein phosphorylation relevant to spine dynamics (Bruning et al., 2019; Cirelli and Tononi, 2020). The circadian clock plays a substantial role in sleep architecture, and global loss of BMAL1 disrupts sleepwake patterns (Laposky et al., 2005; Qiu et al., 2019). Therefore, we carried out continuous 24-hour electroencephalogram/electromyogram (EEG/EMG) recordings to determine baseline sleep-wake patterns in aged *Bmal1* and WT mice over 3 days. A trend towards increased wakefulness was observed in the *Bmal1* cKO mice, (**Fig. 8 A-C**). During the non-active light phase/subjective night, aged *Bmal1* cKO mice spent less time in REM sleep (**Fig. 8 C**; *p*=0.0108) compared to WT mice. Similarly, during the active dark phase/subjective day, aged *Bmal1* cKO mice spent more time awake (**Fig. 8 A**; *p*=0.0366) and less time in REM sleep (**Fig. 8 C**; *p*=0.0253).

**Figure 8.**
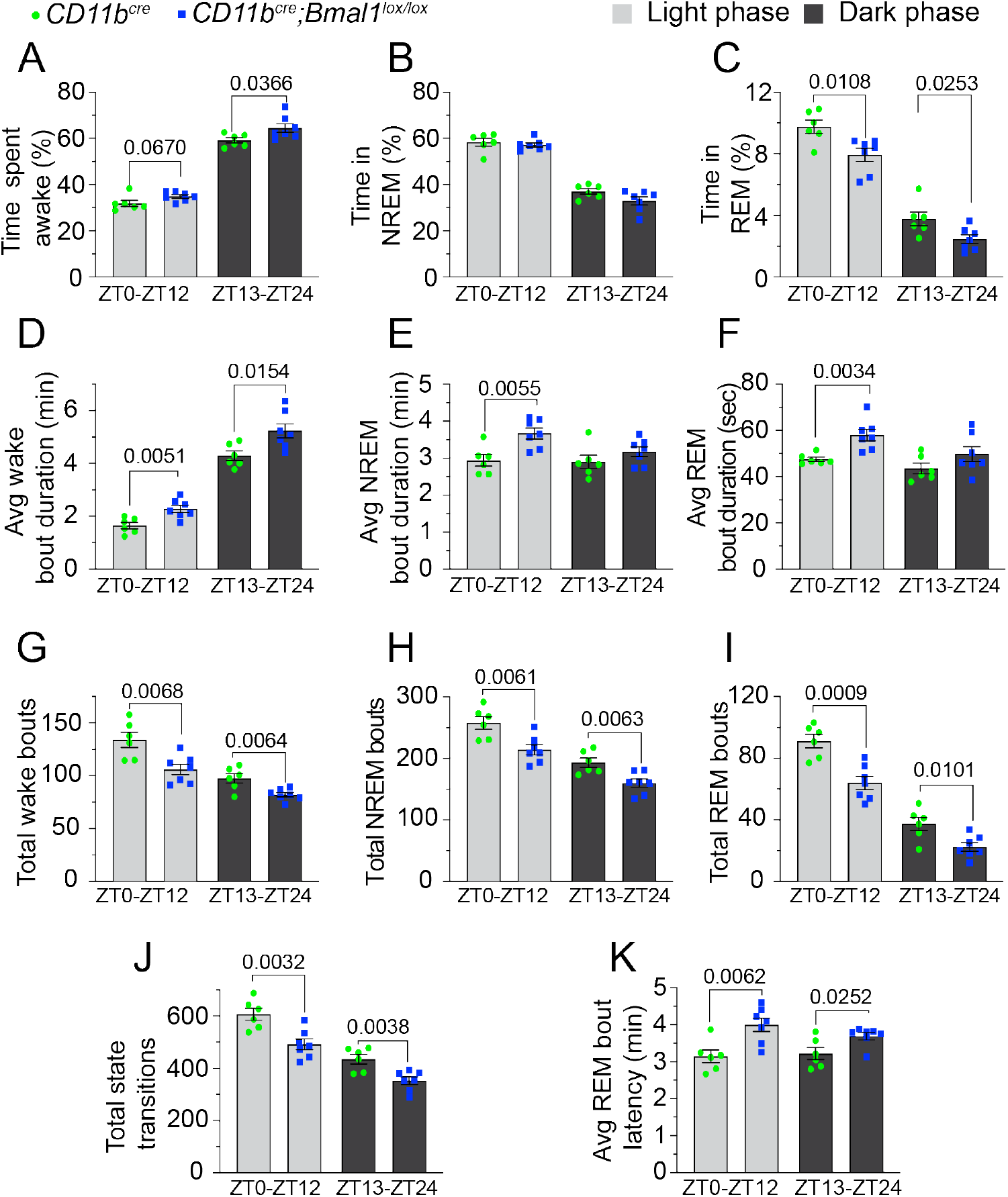
Sleep-wake behavior is disrupted in aged *Bmal1* cKO mice. Baseline sleep-wake behavior was assessed over continuous 24-hr recordings of aged *CD11b^cre^* (18-20 months; n = 6) and *CD11b^cre^;Bmal1^lox/lox^* (n = 7) mice under normal light-dark (12-hr light:12-hr dark) conditions. **(A-C)** Percentage of time spent in wake **(A)**, NREM sleep **(B)**, and REM sleep **(C)** across the entire 12-hr light/dark period. **(D-F)** The average duration of individual wake **(D)**, NREM **(E)**, and REM **(F)** bouts across the entire 12-hr light/dark phase. **(G-J)** The total number of wake **(G)**, NREM **(H)**, and REM **(I)** bouts across the 12-hr light/dark phases. **(J)** Transitions between sleep-wake states across the 12-hr light/dark phases. **(K)** The average REM bout latency during the 12-hr light/dark periods.

During both the light and dark phases, aged *Bmal1* cKO mice exhibited an overall trend towards prolonged sleep-wake cycling. For example, during the non-active light phase/subjective night, the average duration of individual wake (**Fig. 8 D**; *p*=0.0051), NREM (**Fig. 8 E**; *p*=0.0055), and REM (**Fig. 8 F**; *p*=0.0034) bouts was increased, while the total number of wake (**Fig. 8 G**; *p*=0.0068), NREM (**Fig. 8 H**; *p*=0.0061), and REM (**Fig. 8 I**; *p*=0.0009) bouts, and transitions between all sleep-wake states (**Fig. 8 J**; *p*=0.0032) were decreased in the aged *Bmal1* cKO mice compared to WT. Similarly, during the active dark phase/subjective day, the average wake bout duration was increased (**Fig. 8 D**; *p*=0.0154) and the total number of wake bouts (**Fig. 8 G**; *p*=0.0064), NREM bouts (**Fig. 8 H**; *p*=0.0063), REM bouts (**Fig. 8 I**; *p*=0.0101), and state transitions (**Fig. 8 J**; *p*=0.0038) was decreased in the aged *Bmal1* cKO mice. Additionally, the average REM bout latency (time from NREM onset to REM onset) was increased in the aged *Bmal1* cKO mice during both the light (**Fig. 8 K**; *p*=0.0062) and dark (**Fig. 8 K**; *p*=0.0252) phases compared to WT.

Next, we performed an EEG spectral power analysis by averaging the relative values of delta 1 (0.5-2 Hz), delta 2 (2.5-4.5 Hz), theta (5-9 Hz), alpha (6-10 Hz) and beta (15.5-20 Hz) activity during wake, NREM, and REM sleep during the non-active light phase/subjective night. Aged *Bmal1* cKO mice exhibited a relative shift in spectral power characteristics during the wake period, with delta 1 power significantly decreased (*p=0.0084*), and theta power increased (*p=0.0144*), compared to aged WT mice (**Fig. S6 A**). No differences were observed during NREM and REM periods (**Fig. S6 B-C**). In humans, increased delta power and decreased theta power during wake has been correlated with subjective feelings of sleepiness (Bylsma et al., 1994; Painold et al., 2010), and in mice the wake-promoting drug modafinil decreases delta power and increases theta power during wakefulness (Willie et al., 2005; Vas et al., 2020). Taken together, the observed decreased delta and increased theta power, which are signatures of lower sleep pressure, suggests increased wakefulness in the aged *Bmal1* cKO mice. Interestingly, the delta 2 EEG band, which is also associated with homeostatic sleep pressure (Hubbard et al., 2020) was unaffected in the *Bmal1* cKO mice (**Fig. S6**).

These results indicate increased wakefulness during both light and dark periods in aged *Bmal1* cKO mice and suggest a role of circadian-tuned microglia in regulating overall neuronal excitability during the sleep-wake cycle.

### Microglial deficiency of BMAL1 cKO alters response to sleep deprivation

Core clock genes influence compensatory mechanisms induced by sleep deprivation (Naylor et al., 2000; Laposky et al., 2005). Given that the loss of *Bmal1* in microglia in aged mice was sufficient to disrupt the sleep-wake cycle and change sleep architecture, we explored the effects of sleep deprivation in these mice. We subjected aged *Bmal1* cKO and WT mice to a 4-hour sleep deprivation protocol during the non-active light phase/subjective night and then assessed sleep recovery. During the light phase recovery period, a rebound effect was observed in both the WT and *Bmal1* cKO mice, with both groups spending less time awake (**Fig. 9 A and D**; WT: *p=0.0004*; Bmal1 cKO: *p<0.0001*) and more time in NREM sleep (**Fig. 9 B and E**; WT: *p=0.0010*; Bmal1 cKO: *p=0.0004*) compared to their baseline levels from the same timeframe. An increase in the time spent in REM sleep was observed in aged *Bmal1* cKO mice (**Fig. 9 C and F**; *p=0.0445*) compared to WT mice. During the dark phase recovery period, WT mice continued to spend less time awake (**Fig. 9 A and D**; *p<0.0001*) and more time in NREM sleep (**Fig. 9 B and E**; *p=0.0001*) compared to baseline, whereas the amount of time *Bmal1* cKO mice spent in wake, NREM, and REM sleep did not differ from baseline (**Fig. 9 A and F**).

**Figure 9.**
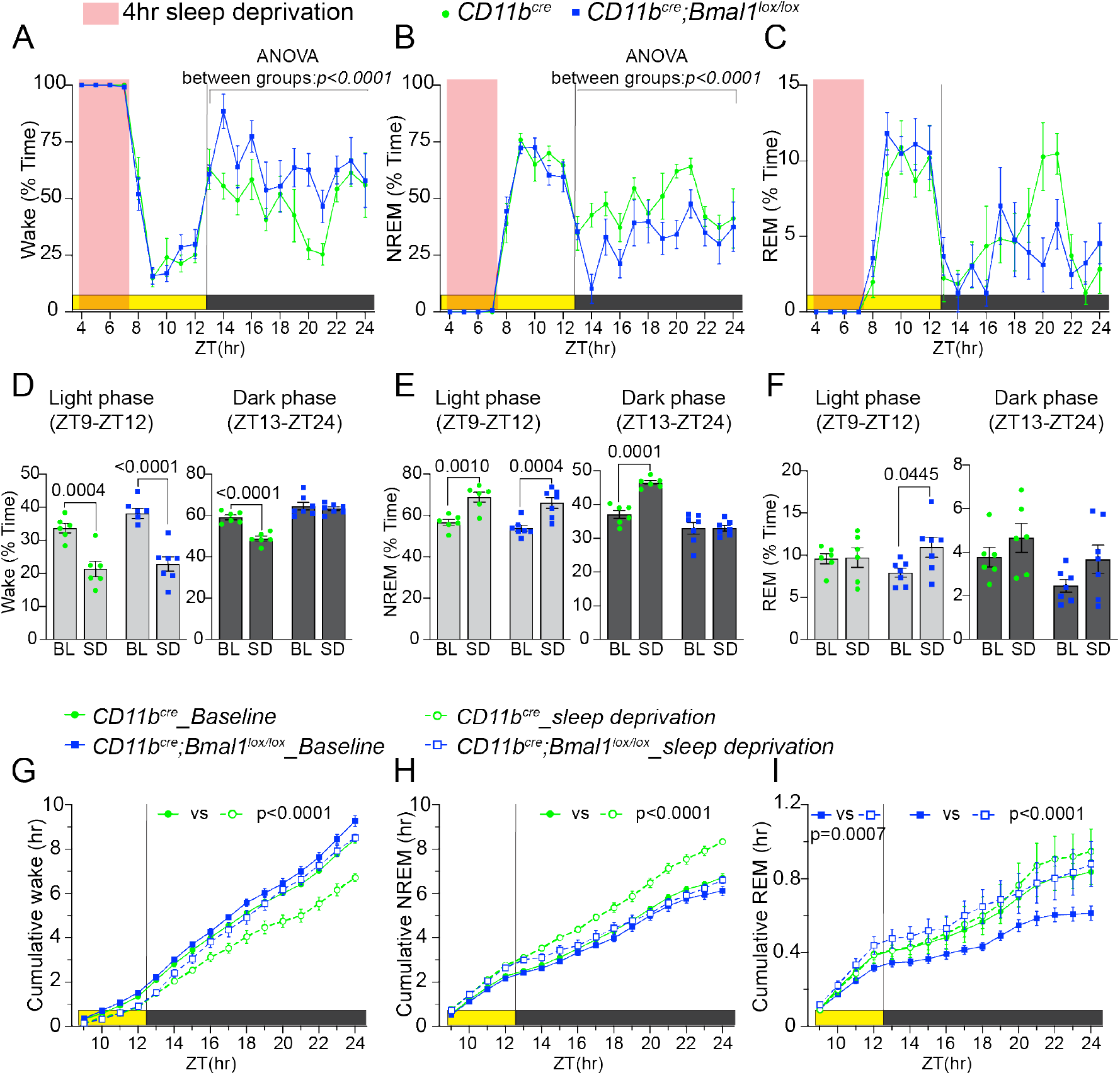
Microglial BMAL1 deficiency alters recovery from sleep deprivation. Data are represented as the mean ± SEM. Significant between group differences in A-B are expressed as *P*-values calculated using Two-way ANOVA. All significant within group differences between baseline (BL) and sleep deprivation (SD) are expressed as adjusted *P*-values generated from Sidak’s post hoc analysis following Two-way ANOVA. **(A-C)** Percent of time per hour spent in wake **(A)** NREM sleep **(B)** and REM sleep **(C)** during a 4-hr sleep deprivation (pink) and subsequent recovery during the remaining light phase and full 12-hr dark phase in aged *CD11b^cre^* (18-20 months; n = 6) and *CD11b^cre^;Bmal1^lox/lox^* (n = 7) mice. **(D-F)** Comparisons of the percentage of time spent in wake **(D)** NREM sleep **(E)** and REM sleep **(F)** across the entire 4-hr light phase recovery period (left) and 12-hr dark phase recovery period (right) during baseline (BL) and sleep deprivation (SD) recordings. **(G-I)** Cumulative amount (total hours) of wake **(G)**, NREM **(H)** and REM **(I)** across the entire recovery period during both BL and SD recordings. Data are represented as the mean ± SEM. *P*-values were calculated using two-tailed Student’s *t*-test. ZT = zeitgeber time.

The cummulative amount of time spent in wake, NREM, and REM sleep did not differ between groups during the light phase recovery period (**Fig. 9 G-H**). However, during the dark phase recovery period, the cummulative time spent by WT mice in wake was decreased (**Fig. 9 G-H**; *p<0.0001*), and increased in NREM (**Fig. 9 G-H**; *p<0.0001*). Unlike WT mice, *Bmal1* cKO mice exhibited no compensatory changes to sleep deprivation in wake and NREM states (**Fig. 9 G-H**). While the cumulative time spent by WT mice in REM did not differ from baseline after sleep deprivation in both the light and dark phase recovery periods, the cumulative time spent in REM was significantly increased from baseline in *Bmal1* cKO mice during the light phase (**Fig. 9 I**; *p=0.0007*) and dark phase (**Fig. 9 I**; *p<0.0001*) recovery periods.

Taken together, these results suggest that the homeostatic response to sleep deprivation is markedly blunted in aged *Bmal1* cKO mice.

## Discussion

The results presented here demonstrate the importance of the clock protein, BMAL1, in microglial function in aging, and highlight the importance of homeostatic circadian interactions between microglia and neurons. Here, we show that with aging, microglial BMAL1 deficiency leads to deficits in non-spatial and spatial learning and memory and increased anxiety. Young *Bmal1* cKO mice performed better than WT mice in spatial learning and memory (Wang et al., 2021) but showed no other differences suggesting that with loss of *Bmal1*, compensatory mechanisms that are activated in young mice are not sustained as the mice age. In aged *Bmal1* cKO mice, electrophysiology of the CA1 hippocampal circuit demonstrated reduction of long-term hippocampal plasticity that was accompanied by a decrease in synaptic vesicle release. These data indicate that synapses in aged *Bmal1* cKO mice may not be functional, an interpretation supported by the observed increase in immature spines by Golgi staining. Additional investigation revealed that microglial BMAL1 deficiency results in reduced deposition of C1q on synapses and reduced microglial synaptic engulfment. This functional deficit may underlie the observed deficits in hippocampal function as well as the additional deficits in the sleepwake cycle and response to sleep deprivation in aged *Bmal1* cKO mice.

Investigation of basal synaptic transmission, short- and long-term synaptic plasticity revealed dysregulated presynaptic vesicle release that reduced the magnitude of long-term hippocampal plasticity in aged *Bmal1* cKO mice. The increased PPR observed in aged *Bmal1* cKO mice suggests reduced availability of pre-synaptic calcium that is required for neurotransmitter release, which would lead to decreases in the readily releasable pool and reduce induction of LTP. Overall the differences in PPR and fEPSP observed between aged *Bmal1* cKO and WT mice indicate a difference in density of functional synapses within the CA1, an interpretation supported by the higher proportion of filopodial-like spines with Golgi stain in aged *Bmal1* cKO mice, and consistent with studies correlating dendritic spine density with neuronal plasticity and memory formation (Mahmmoud et al., 2015; Borczyk et al., 2019). Phosphorylated CAMKII, which facilitates induction of LTP (Lisman et al., 2012; Herring and Nicoll, 2016) by mediating the insertion of endogenous post-synaptic AMPA receptor, did not differ between aged *Bmal1* cKO and WT mice. However, levels of the AMPA receptor subunit, GluA1, which is critical to mediating LTP (Jiang et al., 2021), were significantly decreased in the CA1 hippocampal region of aged *Bmal1* cKO. The reduced density of AMPA receptors at the post-synapse may explain the impaired induction and maintenance of LTP observed in the aged *Bmal1* cKO mice.

Microglia engulf non-functioning dendritic spines and play a critical role in synapse elimination and refinement of neural circuits (Miyamoto et al., 2016; Wang et al., 2020). The observed increase in levels of pre- and post-synaptic proteins, immature synapses by Golgi and the reduction in engulfment of of synaptic components by BMAL1-deficient microglia in aged mice support a critical role for microglial BMAL1 in synaptic pruning. We observed a reduced deposition of C1q at CA1 synapses in aged *Bmal1* cKO hippocampus, where C1q promotes microglial engulfment of synaptic material (Fonseca et al., 2017; Wang et al., 2020) through interaction with the microglial C3 receptor (Perry and O’Connor, 2008; Presumey et al., 2017). The reciprocal increase in C3 may be due to its accumulation in response to decreased C1q and C3 convertase levels. Our results are consistent with another study where global deletion of *Bmal1* increased levels of C3 in the brain, and deletion of REV-ERBα, a negative regulator of BMAL1 expression, increased synaptic phagocytosis (Griffin et al., 2020). Indeed a mutant amyloid precursor protein (APP) transgenic mouse model lacking C1q also demonstrated a similar increase in C3 (Zhou et al., 2008). Additionally, deficiency of the circadian clock protein, CRY1, decreases levels of C1q (Cao et al., 2017), and BMAL1 is known to bind the promoter regions of PU.1 and MafB, two transcription factors that regulate C1q expression (Hatanaka et al., 2010; Oishi et al., 2017; Tran et al., 2017). Therefore, a role for BMAL1 deficiency in disrupting synaptic pruning could be explained by the loss of circadian regulation of complement proteins.

The increase in IBA1, reduced morphological complexity, and decrease in lysosomal proteins CD68, LAMP1 and p62 in aged *Bmal1* cKO mice suggest that the microglia are activated but poorly phagocytic. Transcriptomic analysis was notable for a significant enrichment of genes associated with the omegasome, a transient structure required for autophagosome formation that is dependent on p62 expression (Hsieh and Yang, 2019). One interpretation is that lysosomal dysfunction in BMAL1-deficient microglia results in failure of proper omegasome and autophagosome formation, leading to defective synaptic engulfment, an effect further compounded by the reduced C1q deposition on synapses that need to be pruned.

Microglial molecular processes and structure exhibit a daily rhythmicity that cycles with sleepwake states, with larger soma and increased expression of CD11b, a subunit of the complement receptor, CR3, at the beginning of the sleep cycle, and smaller soma with extensive and motile processes during the wake cycle (Choudhury et al., 2020; Deurveilher et al., 2021). Spine elimination mediated by microglia occurs during sleep, and is associated with increased C1q and C3, phagocytosis, and decreases in synaptic-associated proteins such as PSD95 and synapsin I (Choudhury et al., 2020). There are sleep-dependent increases in C3, C4, and C5 that are abrogated with sleep deprivation (Reis et al., 2011) and potentially function in synapse remodeling. Our results demonstrate a profound effect of microglial deficiency of BMAL1 on the sleep-wake cycle as well as the homeostatic need for sleep after sleep deprivation. Previous studies have demonstrated altered EEG delta power dynamics in response to sleep deprivation from loss of or mutated clock genes (Franken et al., 2006). Global deletion of *Bmal1* in mice has been shown to increase total sleep time and sleep fragmentation (Laposky et al., 2005), however in that study, *Bmal1* was ablated in all cell types including neurons. New learning is encoded during the wake state and sleep consolidates and stabilizes the memory of the learned experience (Tononi and Cirelli, 2014; de Vivo et al., 2017; Diering et al., 2017). With the loss of microglial BMAL1, this process is disrupted and may account for the cognitive deficits and sleepwake deficits observed in aged *Bmal1* cKO mice.

In conclusion, microglial BMAL1 function is critical to maintaining hippocampal plasticity and memory, the sleep-wake cycle, and sleep homeostasis in aging. BMAL1 deficiency in microglia causes profound disruptions in microglial morphology, lysosomal function and generation of C1q, an essential mediator of synaptic engulfment and pruning necessary for the maintenance of neural circuitry.

## Acknowledgements

This work was supported by RF1AG070839 (KIA), 1P30 AG066515 (KIA), American Heart Foundation/Allen Frontiers Award (KIA), The Zhang-Jiang Research Fund (KIA). We are grateful to the Stanford University Cell Sciences Imaging Core Facility (RRID:SCR_017787) for their invaluable support especially Kitty Lee, and to Qian Wang for mouse colony breeding and aging. We are also grateful to Edward N. Wilson and Congcong Wang for review of the manuscript. KIA is a Chan Zuckerberg Biohub investigator.

## Author contributions

C.A. Iweka and K.I. Andreasson conceived and planned the study. C.A. Iweka, M. Cabrera, C.T Pasternak and E. Blacher carried out behavioral, transcriptomic, and immunocytochemical experiments; E. Seigneur and L. deLecea carried out sleep experiments; A.L Hernandez and F.M Longo carried out electrophysiology; S.H Paredes analyzed transcriptomic data. C.A Iweka and K.I wrote the manuscript.

