## Supplementary Materials for "Myeloid deficiency of the intrinsic clock protein BMAL1 accelerates cognitive aging by disrupting microglial synaptic pruning"

### Supplementary Data

I. **Supplementary Figures and legends**

II. **Methods**

III. **References**

### I. SUPPLEMENTARY FIGURES AND LEGENDS

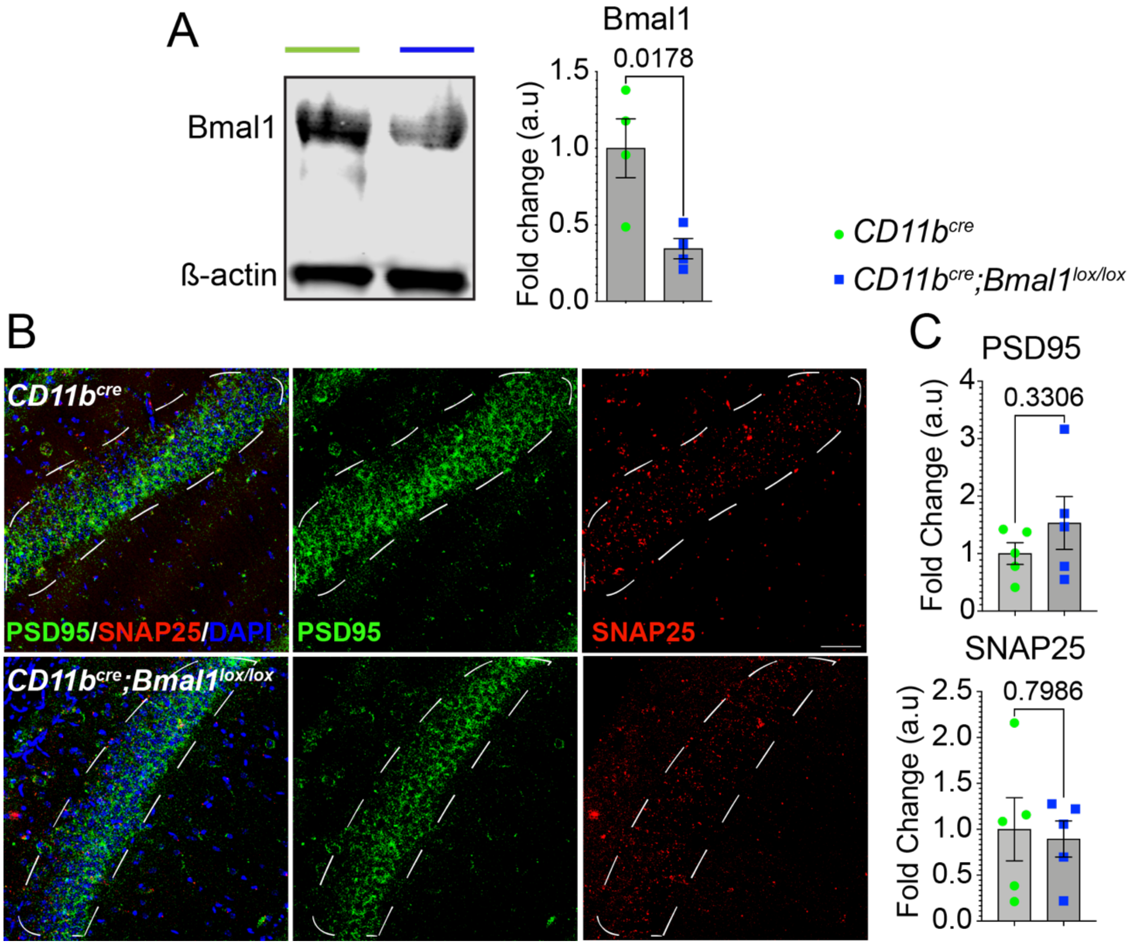


**Figure S1. Young *Bmal1* cKO mice do not demonstrate changes in synaptic density proteins, PSD95, or SNAP25.**

**(A)** Representative immunoblot and quantification of BMAL1 protein levels in peritoneal macrophages show 50% reduction in protein expression in *Bmal1* cKO mice.

**(B)** Representative images of PSD95 (green) and SNAP25 (red) expression in the CA1 hippocampal region of young (3-6 months) *CD11b*$^{cre}$ and *CD11b*$^{cre}$*;Bmal1*$^{lox/lox}$ mice (n = 5/group). Scale bar, 50 μm. White dotted lines in (**B**) indicate region of interest quantified in (**C**).

(**C**) Mean fluorescence intensity of PSD95 and SNAP25 from **(B)**.

Data are represented as the mean ± SEM. P-values were calculated using two-tailed Student's t-test.

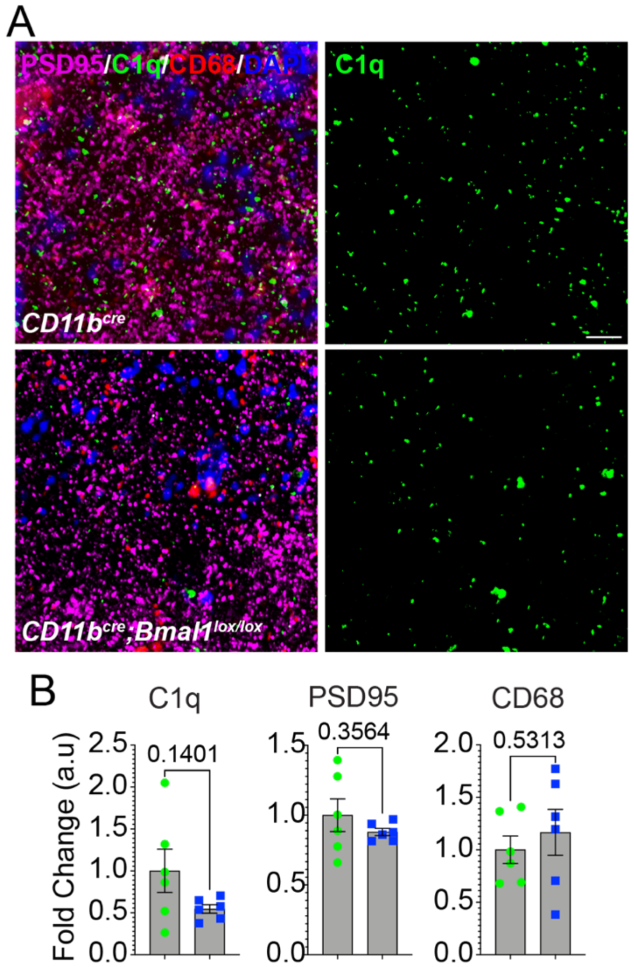


**Figure S2. Young *Bmal1* cKO mice do not demonstrate changes in C1q or CD68 in CA1 hippocampus.**

**(A)** Representative images of PSD95 (magenta) and C1q (green) and CD68 (red) expression in the hippocampal CA1 region in young mice (3-6 months) *CD11b*$^{cre}$ and *CD11b*$^{cre}$*;Bmal1*$^{lox/lox}$ mice (n = 6). Scale bar, 20 µm.

**(B)** Mean fluorescence intensity of C1q, PSD95 and CD68 from **(A).**

Data are represented as the mean ± SEM. P-values were calculated using two-tailed Student's t-test.

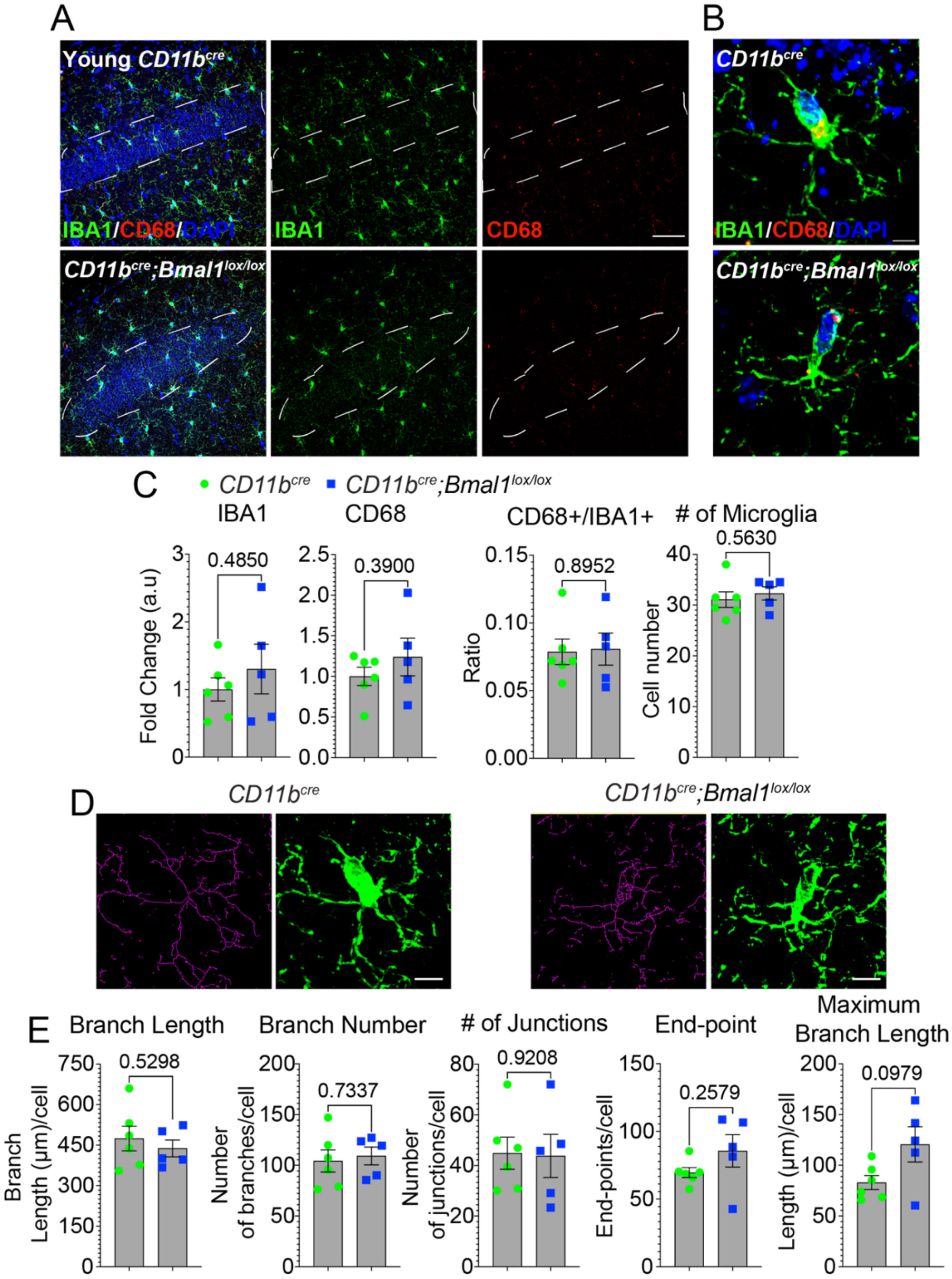


**Figure S3. Young *Bmal1* cKO mice do not demonstrate changes in microglial activation and morphology.**

**(A)** Representative confocal images of IBA1 (green) and CD68 (red) expression in the hippocampal CA1 region in young mice (3-6 months) *CD11b$^{cre}$* (n = 6) and *CD11b$^{cre}$;Bmal1$^{lox/lox}$* (n = 5). Scale bar, 20 µm.

**(B)** Higher magnification, scale bar, 5 µm. White dotted lines in (**A**) indicate region of interest quantified in **(C)**.

(**C**) Mean fluorescence intensity of IBA1, CD68, proportion of IBA1-positive (+) cells that are CD68+, and the number of DAPI+IBA1+ microglia from **(B)**.

**(D)** Representative images of skeletonized microglia overlaid on original image from *CD11b*$^{cre}$ and *CD11b*$^{cre}$*;Bmal1*$^{lox/lox}$ microglia from young mice.

**(E)** Quantification of microglial complexity. Scale bar, 5 μm. Every measurement that contained ≤ 2 endpoints with a maximum branch length of less than the cutoff value of 0.5 μm was removed from the analysis.

Data are represented as the mean ± SEM. *P*-values were calculated using two-tailed Student's *t*-test.

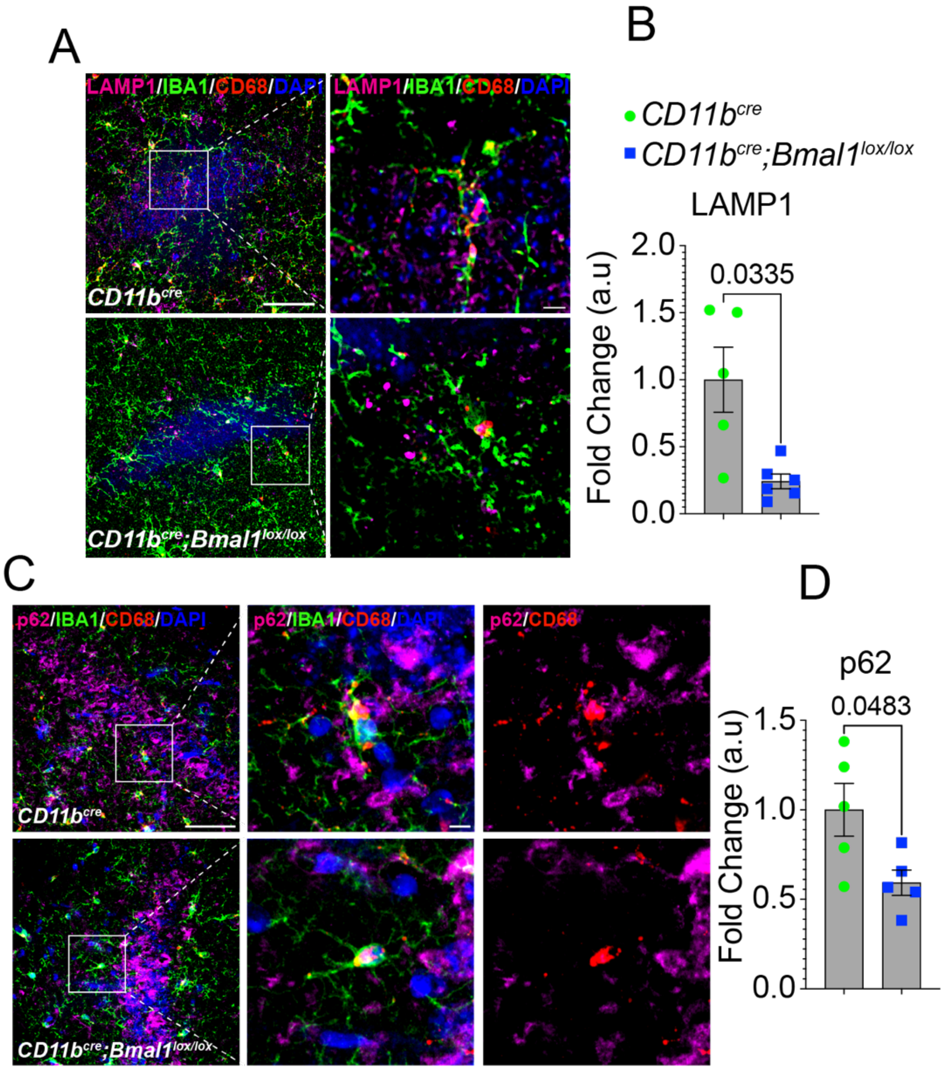


**Figure S4. Microglial BMAL1 deficiency decreases LAMP1 in the CA1 hippocampal region.**

**(A)** Representative confocal images of LAMP1, IBA1 and CD68 expression in the CA1 hippocampal region in aged *CD11b$^{cre}$* (18-20 months; n = 5) and *CD11b$^{cre}$;Bmal1$^{lox/lox}$* (n = 6) mice. Scale bar, 5 μm and 20 μm.

**(B)** Mean fluorescence intensity of LAMP1 in IBA1+ cells.

**(C)** Representative confocal images of p62, IBA1 and CD68 expression in the CA1 hippocampal region of *CD11b$^{cre}$* (n = 5) and *CD11b$^{cre}$;Bmal1$^{lox/lox}$* (n = 5) aged mice (18-20 months). Scale bar, 5 μm and 20 μm.

**(D)** Mean fluorescence intensity of p62 in IBA1+ cells. White arrows show TFEB and p62 localized in CD68+ structures in IBA1+ microglia.

Data are represented as the mean ± SEM. *P*-values were calculated using two-tailed Student's *t*-test.

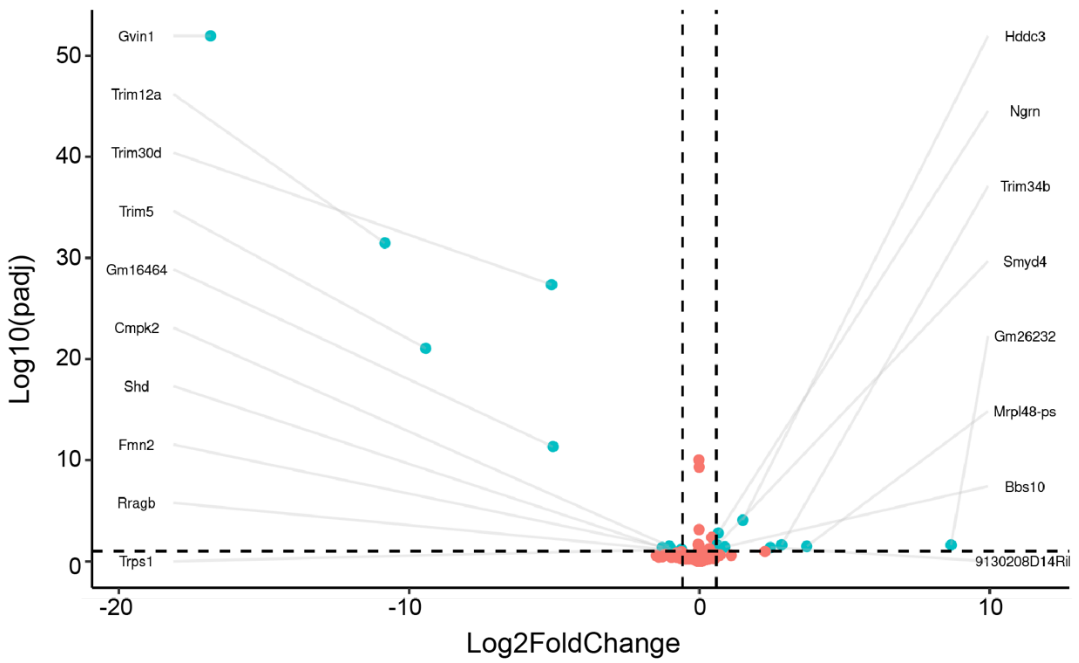


**Figure S5. BMAL1 deficiency in microglial gene expression in young mice.**

Volcano plot showing the -log10 p-adjusted value versus -log2 fold change of normalized counts between young *CD11b*$^{cre}$ and *CD11b*$^{cre}$*;Bmal1*$^{lox/lox}$ mice. Each dot represents a single transcript. Cyan dots denote significant differentially expressed genes. Orange dots denote genes that are unchanged between the genotypes.

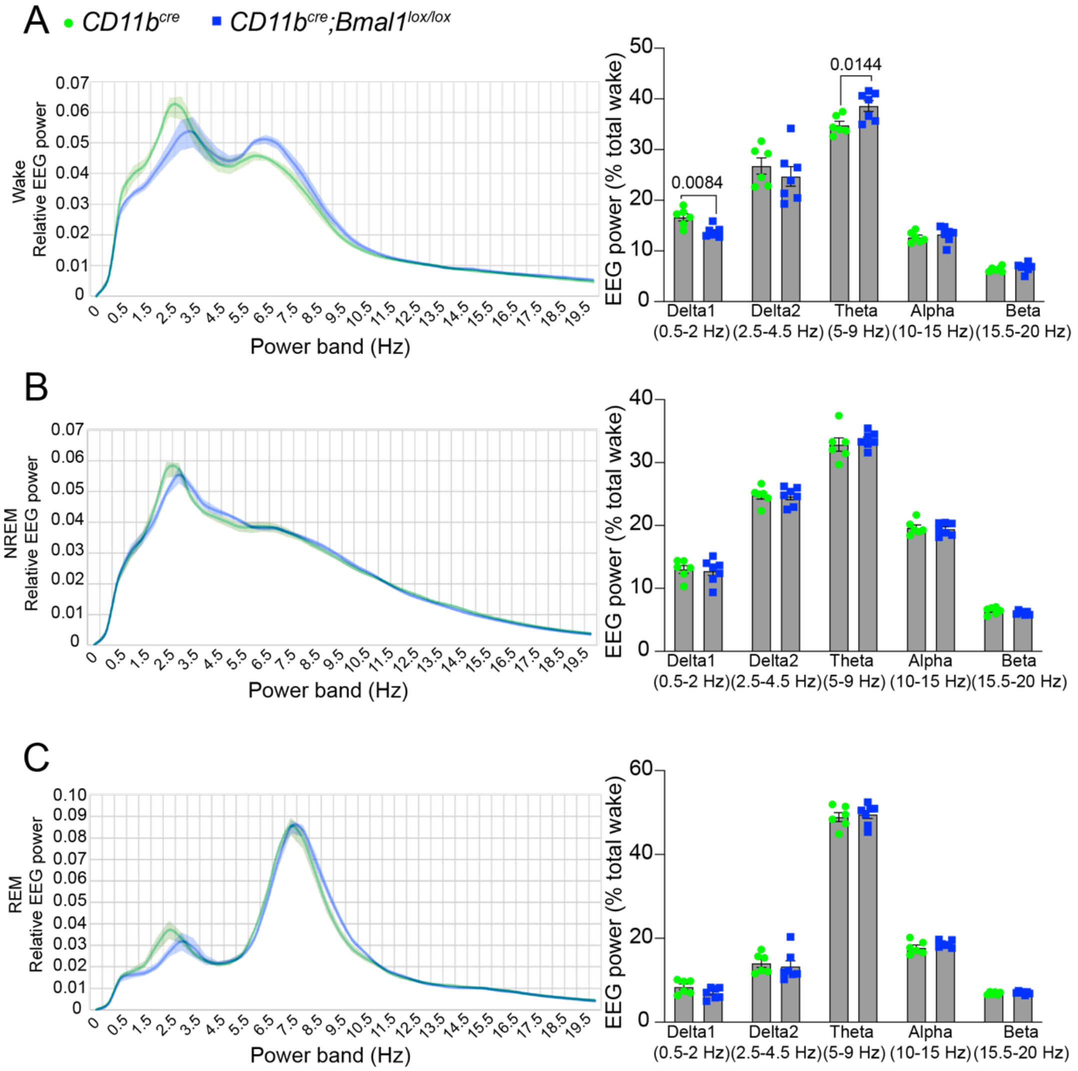


**Figure S6. Baseline EEG spectral power characteristics are shifted during the wake period in aged *Bmal1* cKO mice.**

**(A-C)** The average EEG power densities in the delta 1 (0.5-2 Hz), delta 2 (2.5-4.5 Hz), theta (5-9 Hz), alpha (6-10 Hz) and beta (15.5-20 Hz) frequency bands during wake **(A)** NREM sleep **(B)** and REM sleep **(C)** in *CD11b*$^{cre}$ (n = 6) and *CD11b*$^{cre}$*;Bmal1*$^{lox/lox}$ (n = 7) aged mice (18-20 months). Values were averaged across a 4-hr period during the light phase (ZT8– ZT11). Data are represented as the mean ± SEM. *P*-values were calculated using two-tailed Student's *t*-test.

**Video 1. Decreased engulfment of PSD95 puncta by CD68+ microglia in aged *Bmal1* cKO mice.** Representative 3D image of PSD95 puncta (magenta) within CD68+ lysosomes (red) in IBA1+ microglia (green). Scale bar, 5 µm.

### MATERIALS AND METHODS

#### Animals

All experiments and procedures were performed in accordance with protocols approved by the Institutional Animal Care and Use Committee (IACUC) at the National Institutes of Health. All protocols were approved by the Institutional Animal Care and Use Committee at Stanford University. All animals were housed in an environmentally controlled, pathogen-free barrier facility on a 12-hour light–dark cycle, temperature, and humidity, with food and water available ad libitum. *B6.129S4(Cg)-Arntl$^{tm1Weit}$/J* mice (*Bmal1$^{lox/lox}$*) on a C57BL/6 background were purchased from Jackson laboratory and crossed with *CD11b$^{cre}$* mice to generate myeloid conditional knockouts of *Bmal1* (*Bmal1* cKO). All tissue samples were isolated between ZT2 and ZT4 (Zeitgebers time). This is the time interval where *Bmal1* expression is maximal in different tissues and cell types including myeloid cells (Akashi and Takumi, 2005; Nguyen et al., 2013). For behavioral tests, all animals were placed in the testing room equipped with automated activity monitoring system, VideoTrack (ViewPoint, Stoelting Co, Illinois).

#### Immunofluorescence

Male *CD11b$^{cre}$* and *CD11b$^{cre}$;Bmal1$^{lox/lox}$* littermates were euthanized by $CO_2$ inhalation and then transcardially perfused with 10 mM PBS. Brain tissue was isolated and immersed in 4% paraformaldehyde (PFA; 15714-S; Electron Microscopy Sciences, Hatfield, PA) overnight and serially cryopreserved in 20% sucrose and 30% sucrose. 5 µm brain sections were coronally sectioned using a sliding microtome (Microm HM430, Thermo Fisher Scientific, Waltham, MA). Brain tissue slices were placed in freezing media and stored at -80˚C. For immunostaining, brain tissue samples were washed 3× in PBS for 5 minutes and permeabilized in 0.2% triton-X100 in PBS for 20 minutes at room temperature. Samples were then washed in PBS and incubated for 1 hour in blocking buffer (0.2% triton-X100 supplemented with 10% normal donkey serum in PBS) and then in primary antibody (see Appendix for list of antibodies) overnight at 4˚C. Next, the samples were washed 3× in PBS for 5 minutes and then incubated in secondary antibody (see Appendix for the list of antibodies) for 2 hours. Samples were washed 3× in PBS for 5 minutes and then incubated in Hoechst stain for 10 minutes in PBS. Samples were washed in PBS and then mounted with Prolong$^{TM}$ Gold antifade reagent (P36930; Invitrogen, Eugene, OR) onto Superfrost Plus slides (Thermo Fisher Scientific, Waltham, MA). For each slide, 2 or 3 brain sections were collected, and images acquired on Zeiss 780 LSM confocal microscope (Carl Zeiss, Thornwood, NY) using either 20×/0.45 NA dry objective lens 40×/0.95 NA oil immersion objective lens, or 63×/1.4 NA oil immersion objective lens. For analysis of LAMP1, regions of interest were drawn around IBA1+ cells in the CA1 region of the hippocampus and mean fluorescence intensity determined by ImageJ (Schneider et al., 2012). Morphological analysis of microglia was performed as described (Young and Morrison, 2018).

**Open Field Testing**

The open field task was performed as previously described (Seibenhener and Wooten, 2015). The open field consisted of a plastic box (50cm × 50cm × 50cm) and was illuminated with 150 lux and divided into four quadrants, each with a 25 cm × 25 cm central zone and a 10 cm surrounding border zone. Male $CD11b^{cre}$ and $CD11b^{cre};Bmal1^{lox/lox}$ littermates were placed individually into the periphery of the open field arena and movement and activity of the mice was recorded for 30 minutes. The open field tests were performed during the dark phase. Each arena was cleaned with ethanol between mice. The following motor activity and anxiety-related parameters were recorded: number of entries into either the central or the border zone, the time spent, and the distance traveled in each zone, and the time spent mobile and immobile in each zone. Defecation was counted at the end of the test. All trials were recorded an automated activity monitoring system, VideoTrack (ViewPoint, Stoelting Co, Illinois). All parameters were calculated using Microsoft Excel and analyzed in GraphPad Prism software.

**Novel Object Recognition**

The novel object recognition task was performed as previously described (Leger et al., 2013). $CD11b^{cre}$ and $CD11b^{cre};Bmal1^{lox/lox}$ littermates were habituated to the arena by allowing the mice to roam in the arena for 5 minutes. On day 2, two identical objects were placed in the arena at opposite ends and each mouse was allowed to explore the objects for 10 minutes with a 3-minute interval and then allowed exploration for another 10 minutes. On day 3, one object was replaced with a novel object and each mouse was allowed to explore the arena for 10 minutes. The arena and objects were wiped clean with ethanol between each trial and each mouse. Memory acquisition occurred when there was greater than 50% preference for the new object. All trials were recorded using an automated activity monitoring system, VideoTrack (ViewPoint, Stoelting Co, Illinois). All parameters were calculated using Microsoft Excel and analyzed in GraphPad Prism software.

**Barnes Maze**

Barnes maze task was performed as previously described (Pitts, 2018). Briefly, a large circular platform containing 20 holes on the outer edge was centered over a pedestal and elevated approximately 120 cm above the floor. The escape hole consisted of a PVC elbow joint connector that was similar in texture to the maze, while the other holes were left open to the floor. Distinct visual cues were placed at four equally spaced points around the platform. Two standing bright lights were used to motivate an escape response. The escape hole position remained the same for all days of the task. Male $CD11b^{cre}$

and *CD11b*$^{cre}$*;Bmal1*$^{lox/lox}$ littermates performed three trials per day separated by 10 minutes for 3 consecutive days. Mice were placed in a dark chamber in the center of the arena for 10 seconds to ensure a random starting point and then they were allowed to explore the arena until they entered the escape hole or 3 minutes had passed. For each trial conducted on the same day, the starting location for the mouse was altered relative to the escape hole position. If mice failed to identify the escape hole within 3 minutes, they were gently guided by light tapping/directing toward the escape hole and given a score of 3 minutes. Between each trial, both the maze and escape hole were thoroughly cleaned with ethanol to remove any scent cues that might affect performance in subsequent trials. Primary latency, or the time to initial contact with the escape hole, was used as a measure of spatial memory. All trials were recorded an automated activity monitoring system, VideoTrack (ViewPoint, Stoelting Co, Illinois). All parameters were calculated using Microsoft Excel and analyzed in GraphPad Prism software.

**Synaptosomal preparation**

Hippocampal brain tissue isolated from male *CD11b*$^{cre}$ and *CD11b*$^{cre}$*;Bmal1*$^{lox/lox}$ littermates was homogenized in 32 M sucrose in HEPES buffer (145 mM NaCl, 5 mM KCl, 2 mM $CaCl_2$, 1 mM $MgCl_2$, 5 mM glucose, 5 mM HEPES, pH 7.4) and cleared by centrifugation at 600 ×g for 10 minutes at 4˚C. Supernatant was diluted 1:1 in 1.3 M sucrose in HEPES buffer and centrifuged at 20,000 ×g for 30 minutes at 4˚C. Supernatant was saved and stored at -80˚C for comparison. The pellet was resuspended in RIPA lysis buffer (89901; Thermo Fisher Scientific, Waltham, MA) supplemented with protease and phosphatase inhibitors (A32959; Thermo Fisher Scientific, Waltham, MA).

**Western blotting**

Cell lysates or hippocampal synaptosomes were prepared in RIPA lysis buffer and clarified by centrifugation. Proteins from supernatants were quantified using the BCA method with the Pierce™ BCA Protein Assay Kit (23227; Thermo Fisher Scientific, Waltham, MA). Equal concentrations of proteins (20 µg/lane) were separated on SDS-PAGE and transferred onto a polyvinylidene difluoride (PVDF) membrane (1620177; Bio-Rad Laboratories, Inc. Hercules, CA). The membrane was blocked for 1hr with 5% nonfat dried milk in 0.1% Tween20 in 10 mM Tris–HCl pH 7.5 (TBST, 15567027; Gibco, Waltham, MA) or blocked in 5% BSA (bovine serum albumin, A4161; Sigma, St. Louis, MO) in TBST. Membranes were incubated with primary antibodies overnight at 4˚C, washed 3× in TBST for 10 minutes and subsequently incubated with secondary antibody for 2 hours at room temperature. Membranes were washed in 6× in TBST for 5 minutes and blots were imaged on the LiCor Odyssey CLx (LI-COR, Lincoln, NE) and analysis was completed using the associated software.

**Golgi preparation**

Male *CD11b*$^{cre}$ and *CD11b*$^{cre}$*;Bmal1*$^{lox/lox}$ littermates were euthanized by cervical dislocation. Brain tissue was rapidly removed from each mouse and rinsed with distilled water and then transferred into the impregnation solution as specified by the Hito Golgi-Cox OptimStain™ Kit (Hitobiotec Corp. Kingsport, TN). Samples were stored in the dark for 24 hours and then placed in fresh impregnation solution for 2 weeks in the dark at room temperature. Samples were then transferred to a sucrose solution and stored at 4˚C for 12 hours in the dark and then replaced in fresh sucrose solution and stored at 4˚C for 72 hours. Samples were then frozen in Tissue-Tek® O.C.T. compound (Sakura Finetek USA, Inc. Torrance, CA) and then cut into 80 µm sections and allowed to dry. Samples were then rinsed in distilled water twice for 3 minutes and then stained with 20% ammonia solution and 1% sodium thiosulfate solution for 10 minutes and rinsed in distilled water twice for 4 minutes. Samples were serially dehydrated in 50%, 75%, 95% and 100% ethanol, cleared in xylene and coverslips applied using Permount mounting medium (Thermo Fisher Scientific, Waltham, MA). Brightfield z-stack images were acquired on Zeiss 780 LSM confocal microscope (Carl Zeiss, Thornwood, NY) using either 20×/0.45 NA dry objective lens or 63×/1.4 NA oil immersion objective lens.

Apical secondary dendritic segments from layer IV/V cortical pyramidal cells and pyramidal cells in the CA1 hippocampal region that were clearly filled, without breaks or overlaps, and without precipitate, approximately 10 – 20 µm long were analyzed by an observer blinded to genotype to obtain dendritic spine density.

**Microglial isolation for RNA-seq**

Brain tissues from male *Cd11b*$^{cre}$ and *Cd11b*$^{cre}$;*Bmal1*$^{lox/lox}$ littermates were homogenized in Dounce buffer (15 mM HEPES buffer, 0.5% glucose, in HBSS without phenol red), supplemented with RNAase inhibitor (RNAsin, 3335402001, 2000U; Sigma, St. Louis, MO) and then passed through a 70 µm cell strainer (352350; Falcon, Fisher Scientific, Pittsburgh, PA). Cells were pelleted at 400 ×g for 5 minutes at 4˚C and resuspended in MACS buffer (0.5% BSA, 2 mM EDTA in 10 mM PBS). Cells were incubated with myelin removal beads (130-096-433; Miltenyi Biotec, Auburn, CA) at 4˚C for 15 min and then washed in 2 ml MACS buffer supplemented with RNAase inhibitor. Cells were then run through the Miltenyi LD column placed in the MACS magnet (130-043-901 LD Columns; 130-090-976 QuadroMACS Separator; Miltenyi Biotec, Auburn, CA) and columns were subsequently rinsed twice with MACS buffer. Cells were pelleted at 300 ×g for 5 min at 4˚C, resuspended in MACS buffer and incubated with CD11b microbeads (130-093-634; Miltenyi Biotec, Auburn, CA) for 15 min at 4˚C. Cells were washed in MACS buffer at 300 ×g for 10 min at 4˚C and resuspended in MACS Buffer. Cells were separated by Miltenyi LS columns (130-042-401; Miltenyi Biotec, Auburn, CA). Cells were then centrifuged at 300 ×g for 10 min at 4˚C and resuspended in TRIzol LS (10296-028, ThermoFisher, Waltham, MA).

**RNAseq Library construction and sequencing**

A total amount of 1 µg RNA per sample was used as input material for RNA sample preparation. Lack of RNA degradation was confirmed on 1% agarose gels. RNA purity was checked using the NanoPhotometer® spectrophotometer (IMPLEN, CA, USA). RNA integrity and quantification were assessed using the RNA Nano 6000 Assay Kit of the Bioanalyzer 2100 system (Agilent Technologies, CA, USA). Sequencing libraries were generated using NEBNext® UltraTM RNA Library Prep Kit for Illumina® (NEB, USA) following manufacturer's recommendations and index codes were added to attribute sequences to each sample. Briefly, mRNA was purified from total RNA using poly-T oligo-attached magnetic beads. Fragmentation was carried out using divalent cations under elevated temperature in NEBNext First Strand Synthesis Reaction Buffer (5X). First strand cDNA was synthesized using random hexamer primer and M-MuLV Reverse Transcriptase (RNase H-). Second strand cDNA synthesis was performed using DNA Polymerase I and RNase H. Remaining overhangs were converted into blunt ends via exonuclease/polymerase activities. After adenylation of 3' ends of DNA fragments, NEBNext adaptors with hairpin loop structure were ligated to prepare for hybridization. To select cDNA fragments of 150~200 bp in length, library fragments were purified with AMPure XP system (Beckman Coulter, Beverly, USA). Then 3 µl USER Enzyme (NEB, USA) was used with size-selected, adaptor-ligated cDNA at 37 °C for 15 min followed by 5 min at 95 °C before PCR. Then PCR was performed with Phusion High-Fidelity DNA polymerase, Universal PCR primers and Index (X) Primer. Finally, PCR products were purified (AMPure XP system) and library quality was assessed on the Agilent Bioanalyzer 2100 system.

**RNAseq analysis**

Raw mRNA sequencing data was processed with the nf-core/rnaseq pipeline (v3.8.1) (Ewels et al., 2020). Default parameters were used except for removing reads derived from ribosomal RNA (rRNA). Transcript integrity numbers were calculated, and reads were mapped to the mouse reference genome. According to default parameters, reads were mapped with STAR (v2.7.10a)(Dobin et al., 2013) and quantified transcript abundance with Salmon (v1.5.2) (Patro et al., 2017). All output is available upon request. Differential expression analysis (DEA) was performed with DESeq2 (v1.32.0)(Love et al., 2014). Two outlier samples were removed from downstream analysis, but their sequence data is available. Only transcripts with more than five reads in at least each of 5 different samples were considered for DEA. A single four level variable (cKO_Aged, cKO_Young, WT_Aged, and WT_Young) was used to encode the experimental design, and DEA was performed between each pair of conditions with default DESeq2 parameters followed by log fold-change shrinkage with the "apeglm" method (Zhu et al., 2019). Genes were deemed differentially expressed if their fold-change in any direction was greater than 1.5, and their adjusted p-value (Benjamini-Hochberg method) was less than 0.1. Principal component analysis (PCA) was performed on the variance stabilizing transformation of the transcript count matrix within the DESeq2 package as well. Functional enrichment analysis was performed via PANTHER statistical overrepresentation test (Release date: 2022-07-12) (Thomas et al., 2022) on the gene ontology (GO) annotations of the mouse genome (Release date: 2022-03-22). Only genes tested were used as background. GO terms with at least three genes and an adjusted p-value (Benjamini-Hochberg method) of less than 0.1 were considered enriched. Plots for all analysis were created with ggplot2 (Wickham, 2009). All code is available upon request. GEO accession "GSE214514".

**Electrophysiology**

Measurement of hippocampal plasticity was carried out as previously described (Minhas et al., 2021). Male $CD11b^{cre}$ and $CD11b^{cre};Bmal1^{lox/lox}$ littermates were euthanized by cervical dislocation, and the hippocampus was rapidly dissected out into ice-cold (4°C) artificial cerebrospinal fluid (ACSF), saturated with carbogen (95% $O_2$/5% $CO_2$). ACSF consisted of 124 mM NaCl, 4.9 mM KCl, 24.6 mM NaHCO3, 1.20 mM $KH_2PO_4$, 2.0 mM $CaCl_2$, 2.0 mM $MgSO_4$ and 10.0 mM glucose, and pH 7.4. Transverse hippocampal slices, 350 μm thick, were prepared from the dorsal area with the McIlwain tissue chopper (Stoelting, Wood Dale, IL) and transferred to a recovery chamber for at least 1.5 hours with oxygenated ACSF at room temperature before being placed into a submerged-type chamber at 32 °C. Slices were continuously perfused with ACSF at a flow-rate of 1.5 ml/min. Slices were then carefully positioned on a R6501A multi-electrode array (Alpha MED Scientific, Osaka, Japan) with electrodes arrayed in an 8 × 2 matrix with an interpolar distance of 150 μm and each matrix measured at 50 μm × 50 μm. After a 30-minute incubation, fEPSPs in the CA1 hippocampal region were recorded by stimulating downstream electrodes in the CA1 and CA3 regions along the Schaffer collateral pathway. Signals were acquired using the MED64 System (AlphaMED Sciences, Osaka, Japan). The time course of the fEPSP was calculated as the descending slope function for all the experiments. Input/output curves were established by applying increasing stimulus currents to the pathway starting from 10 μA to 90 μA (in 5 μA increments) and recording evoked responses. After input/output curves had been established, the stimulation strength was adjusted to elicit a fEPSP slope at 35% maximal value, which was maintained throughout the experiment. During the baseline recording, a single response was evoked at a 30-second interval for at least 20 minutes. To induce a strong form of long-term potentiation, three episodes of theta-burst stimulation (TBS) were used, each TBS consisting of 10 bursts of 4 stimuli at 100 Hz separated by 200 ms (double pulse-width), followed by a recording of evoked responses beginning 1 minute after the induction of long-term potentiation and continuing every 30 seconds until the end of the experiment. WT control and cKO mice were randomized during the experiment. The mean baseline fEPSP value was calculated and the percentage change from baseline after TBS was analyzed for long-term potentiation.

Paired-pulse ratio (PPR) experiments were conducted as described (Latif-Hernandez et al., 2020). PPR responses to two impulses given at an interval of 10, 20, 50,100, 200, or 500 ms were recorded. During baseline recording, 3 single stimuli (0.1 ms pulse width; 10 second intervals) were measured every 5 minutes and averaged for the 60-minute fEPSP values.

**Surgery for sleep deprivation studies**

Male *CD11b$^{cre}$* and *CD11b$^{cre}$;Bmal1$^{lox/lox}$* littermates were anesthetized with an intraperitoneal (IP) injection of ketamine (100 mg per kg bodyweight) and dexmedetomidine (0.5 mg per kg bodyweight) and placed on a small animal stereotaxic frame (David Kopf Instruments). Mice were then implanted with a custom-made EEG/EMG implant comprised of two stainless steel screws implanted over the right frontal cortex (1 mm anterior to Bregma and 1.5 mm lateral to the midline) and right parietal cortex (1 mm anterior to lambda and 1.5 mm lateral to the midline) for recording EEG, and two stainless steel wires implanted underneath the neck muscles for recording EMG signals. Following the procedure, mice received an IP injection of atipamezole (5 mg per kg bodyweight) to reverse the anesthetizing effects of dexmedetomidine. Following surgery, the mice were individually housed in custom recording chambers/cages and allowed to recover for 2 weeks, followed by an additional 7 days of acclimation to a flexible EEG/EMG connection cable.

**Polysomnographic recordings and analysis**

Male *CD11b$^{cre}$* and *CD11b$^{cre}$;Bmal1$^{lox/lox}$* littermates were housed in reversed 12-hr light/dark cycle, beginning at 7:00 AM (Zeitgeber time (ZT) 0). Three continuous 24-hr recordings were collected beginning at ZT 0, and the average of the 3 recordings was analyzed. The EEG/EMG signals were amplified (Grass Instruments) and digitized at 256 Hz using Vital Recorder (Kissei Comtec America) software. The signals were then digitally filtered (EEG: 0.5-30 Hz, EMG: 10-40 Hz) and spectrally analyzed by fast Fourier transformation, and manually scored in 4-s epochs as wake, non-REM (NREM) sleep, or REM sleep using SleepSign (Kissei Comtec America). We defined wake, NREM sleep, and REM sleep as the following: Wake: desynchronized low-amplitude EEG and heightened tonic EMG activity with phasic bursts; NREM sleep: synchronized, high-amplitude, low-frequency (0.5 – 4.5 Hz) EEG with little to no EMG activity; and REM sleep: desynchronized low-amplitude EEG with a pronounced theta rhythm (5 – 9 Hz) and a flat EMG. For calculating sleep-wake bout numbers and durations, we defined the sleep cycle as beginning at the onset of NREM sleep and ending at the offset of REM sleep or the onset of a wake episode lasting ≥12 seconds, allowing for brief awakenings (<12 seconds) in either case.

For the sleep deprivation and recovery study, the mice were sleep deprived for 4-hours starting at ZT 4 by placing the animals in a novel environment enriched with novel objects (various plastic toys) and novel foods and lacking any bedding material. The live EEG/EMG recordings were monitored and whenever a mouse entered NREM sleep, they were aroused by gentle stroking with a paintbrush. After 4 hours, the mice were returned to their home cage and sleep/wake behavior during the remaining light phase and subsequent dark phase was monitored and analyzed.

**Statistical analysis**

Analyses were performed using GraphPad Prism software version 9 (GraphPad Software, LaJolla, CA). Unless otherwise specified, values represent the means ± SEM with $P < 0.05$ considered statistically significant. Data were analyzed by paired and unpaired Student's *t*-tests, one-way ANOVA, two-way ANOVA, or repeated measures two-way ANOVA with Tukey's posthoc or Bonferroni multiple comparison tests to determine significance. Normality of the distribution of the data was tested with Kolmogorov–Smirnov normality tests using the column statistics function of GraphPad Software.

**Online supplemental materials**

Fig. S1 shows a representative immunoblot of BMAL1 protein in peritoneal macrophages and a representative immunofluorescent image and quantification of PSD95 and SNAP25 expression in the CA1 hippocampal region in young mice. Fig. S2 shows representative immunofluorescent images and quantification of PSD95, C1q and CD68 in the hippocampal CA1 region in young mice. Fig S3 shows representative immunofluorescent images and quantification of IBA and CD68 in the CA1 hippocampal region and the morphological analysis of IBA1+ microglia from young mice. Fig. S4 shows the representative immunofluorescent images and quantification of lysosomal protein LAMP1, and p62 in aged mice. Fig. S5 shows the volcano plot of differentially expressed genes in microglia isolated from young and aged mice. Fig S6 shows the EEG spectral power analysis in aged mice.
